# Predictive Modelling of The Dynamic Patterns of Thinking in Attention-Deficit/Hyperactivity Disorder: Diagnostic Accuracy of Spatiotemporal Fractal Measures

**DOI:** 10.1101/420513

**Authors:** F. Labra-Spröhnle, G. Smith, H. Ahammer, C. Postlethwaite, I. Liu, P. Teesdale-Spittle, M. Frean

## Abstract

**Background:** Attention-Deficit/Hyperactivity Disorder (ADHD) is a neurodevelopmental condition characterized by executive function (EF) dynamics disturbances. Notwithstanding, current advances in translational neuroscience, no ADHD objective, clinically useful, diagnostic marker is available to date.

**Objectives:** Using a customized definition of EF and a new clinical paradigm, we performed a prospective diagnostic accuracy trial to assess the diagnostic value of several fractal measures from the thinking processes or inferences in a cohort of ADHD children and typically developing controls.

**Method:** We included children from age five to twelve diagnosed with a reference standard based on case history, physical and neurological examination, Conners 3^rd^ Edition, and DSM-V™. The index test consisted of a computer-based inference task with a set of eight different instances of the “Battleships” game to be solved. A consecutive series of 18 cases and 18 controls (n = 36) recruited at the primary paediatrics service from the Nelson Marlborough Health in New Zealand underwent the reference standard and the index test. Several fractal measures were obtained from the inference task to produce supervised classification models.

**Results:** Notably, the summarized logistic regression’s predicted probabilities from the eight games played by each children yielded a 100% classification accuracy, sensitivity and specificity in both a training and an independent testing/validating cohort.

**Conclusions:** From a translational vantage point the expeditious method and the robust results make this technique a promising candidate to develop a screening, diagnostic and monitoring system for ADHD, and may serve to assess other EF disturbances.

## 1 Introduction

### 1.1 Scientific and Clinical Background

Attention-Deficit/Hyperactivity Disorder (ADHD) is widely recognised as one of the most common childhood neuro-developmental disorders (Sayal et al., 2018; Bölte et al., 2018). The American Psychiatric Association characterised this condition by chronic, age-inappropriate levels of hyperactivity, inattention, and impulsivity and distinguished three main presentations (2013):

1. Combined presentation: when six or more symptoms of both inattention and hyperactivity/impulsivity have been observed for at least six months.
2. Predominantly inattentive presentation: if six or more symptoms of inattention, but less than six hyperactivity/impulsivity symptoms have persisted for at least six months.
3. Predominantly hyperactive/impulsive presentation: if six or more hyperactivity/impulsivity symptoms, but less than six inattention symptoms have persisted for at least six months.

A recent systematic review and meta-analysis from prevalence studies, estimated that the overall pooled prevalence rate of ADHD is around 7.2% (CI 95%: 6.7 to 7.8) (Thomas et al., 2015). The exact aetiology of ADHD remains unknown and controversial. Moreover its progression is unpredictable and its treatment is complicated and of a limited success (Armstrong and Lezak, 2012). Nonetheless, several genetic, non-genetic and epigenetic interactions have been identified (Hamza et al., 2017; Bhat et al., 2017). ADHD is a major diagnostic challenge for clinicians; no useful bio-markers or “gold standard” diagnostic tests are available to date (Scassellati et al., 2012; Thome et al., 2012; Rothenberger et al., 2015; Jaffee, 2018).

The current diagnosis of ADHD is performed in a subjective manner by collecting observations of symptoms from parents, teachers and clinicians. Notwithstanding, it has been estimated that around 20% of those diagnoses could be mistaken (Elder, 2010; Merten et al., 2017), and hence in recent decades a large amount of translational research in medicine has been carried out trying to devise an objective technology for ADHD diagnosis. Many of the most salient and promising efforts to identify cognitive biomarkers for ADHD rest on the assessment of a controversial construct that has been labelled executive function (EF).

The main reason for targeting EF as a potential source of knowledge to be translated into a diagnostic tool, is that children with ADHD have revealed a pattern of cognitive deficits consistent with pre-frontal EF deficits, namely inattention, difficulty with self-regulation, response inhibition deficits (impulsivity), restlessness or hyperactivity, or in some cases apathy (Inagaki, 2011). Despite the controversy and the more than 30 definitions of the concept of EF (Goldstein and Naglieri, 2013), the currently accepted view refers to a wide set of more or less independent higher order cognitive processes and abilities. These include: self-regulation, reasoning and problem-solving, anticipating, setting goals, planning and decision-making, the ability to sustain attention and resistance to interference, utilisation of feed-forward, feedback and multitasking, cognitive flexibility and the ability to deal with novelty(Chan et al., 2008; Zelazo et al., 1997; Zelazo, P. D., Carlson, S. M., & Kesek, A., n.d.).

The poorly defined boundaries and the lack of a unified and integrative definition of the concept of EF makes initial translational steps very hard to accomplish. Nevertheless, researchers have managed to provide different procedures to assess this construct in experimental and clinical settings (Goldstein and Naglieri, 2013). Most of the outcomes from these procedures and techniques have been deficient and of limited success (Hall et al., 2016; Lange et al., 2014). It seems that the dual issues of (i) a lack of a proper definition and (ii) operationalization of the EF concept present a challenge to the use of measures of EF for the ADHD diagnosis at the individual level that is unsurmontable (Faraone et al., 2014). For example, none of the existing theoretical constructs and their associated procedures to evaluate EF could account for the heterogeneity of ADHD clinical presentation. As a consequence they have never produced acceptable levels of sensitivity and specificity in any ADHD diagnostic accuracy study (Thome et al., 2012).

The concept of EF, although not the term itself, originated in the former Soviet Union. In its original format, this concept integrated neurophysiological, psychological and socio-historical dimensions (Labra-Spröhnle, 2015, 2016a). This notion was born rooted in Anokhin’s theory of “Functional Systems” (Anokhin, 1974), Bernshtein’s “Physiology of Activity” (Bernstein, 1967), Vygotsky’s “Socio-Historical” approach to higher cognitive functions (Vygotsky, 1965) and Filimonov’s principle of graded and pluripotential localisation of functions in the brain (Luria, 1966, 1976, 1980). This conceptual synthesis was used by Luria to analyse the disturbances of higher cortical functions produced by local brain lesions. Luria’s work on frontal lobe and dysexecutive syndrome was crucial to differentiate EF from other aspects of cognitive processes (Luria, 1976, 1980). During the 80s the cognitive revolution took over this construct, translating the original notion into a computational, information processing and modular paradigm (Shallice, 1982; Duncan, 1986; Welsh and Pennington, 1988). As a result of this translation the concept was fragmented and detached from its original sources. Today the cognitive, modular and computational paradigm has been the dominant perspective that frames most of the EF views (cf. Shallice et al.,2018). Moreover this frame is the “official doctrine” that determines how this construct is currently understood and assessed for ADHD diagnosis. Consequently, most of the practical methods for assessing EF for an ADHD diagnosis under the cognitive paradigm rest on measures of performance or achievement (Goldstein and Naglieri, 2013; Griffin et al., 2015; Hoskyn et al., 2017) of the individuals from different particular cognitive domains and are based in open loop tasks (Marken, 1988, 2009). To the best of our knowledge no integrative measures have been produced that could account for the mechanisms and the regulatory aspect of EF across the different cognitive domains and using a “closed-loop task”.

When looking for a common feature that could subserve all the diverse elements, behaviours or cognitive operations that has been put under the umbrella of the EF concept, the first point to notice, is that all of them are manifestations of goal-directed behaviours. The second is that all these goal-directed activities are powered and driven by inferential processes (in a wide sense, i.e., at presentational and representational levels). Inferences are dynamical coordinators that create meaningful implications inside and across different cognitive domains (perceptions, actions, functions, operations) Piaget (1985); Piaget et al. (2013). By these means, inferences regulate and drive the organism-environment system from the very basic sensorimotor levels, as in unconscious innate reflex actions; right up to the highest goal-directed conscious cognitive activities (Peirce, 1998; Piaget et al., 2013; Labra-Spröhnle, 2016a). The fundamental role of the inferential processes (and their subordinated anticipatory behaviours) in the structure of any goal directed behaviour is paramount in the theory of functional systems (Anokhin, 1974) and in the theory of inferences from Peirce (1998), and Piaget (2013). The convergence of these theories inspired one of us to develop in detail an “intensional” definition of EF (Labra-Spröhnle, 2016a). The key implications of this definition for the current work are that EF can be viewed as a cycle of inferential processes that are central to behaviour and cognition. Besides, differences in EF between individuals, including EF disorders, could be revealed by assessing the spatio-temporal structure of inferential dynamics.

Evaluation of behavioural self-regulation (Barkley, 2011, 2012), action selection and decision making (Matthies et al., 2012a; Schepman et al., 2012a,b), shows consistently that goal-directed behaviours are the cardinal features most impaired in in ADHD individuals (Guevara and Stein, 2001). In spite of the fact that inferential processes are at the core of EF, represented in decision making processes, self-regulation, setting goals and strategy selection (Chevalier, 2015; Chevalier et al., 2017; Blair and Ursache, 2011; Sella et al., 2012; Mowinckel et al., 2015), those processes have not previously been directly targeted for EF assessment in ADHD patients.

We suggest here that there is a gap in knowledge at the nexus of EF, inferential processes, and ADHD, caused by the lack of a theoretical synthesis that could integrate different domains of knowledge and the clinical facts associated to these concepts. This circumstance has led to a two-fold methodological deficiency: firstly the absence of an empirical model of human inferential processes, and secondly the absence of an experimental paradigm to assess them. From a translational point of view, these limitations has been hindering the conversion of basic knowledge into a clinical technology for the diagnosis and management of EF disorders such as ADHD.

To overcome this situation, a new approach is advocated herein to tackle EF assessment and the diagnosis of ADHD. The core of this proposal rests in a new conceptualisation and operationalization of EF that permits the mathematical description and classification of the dynamics of human inferential processes (Labra-Spröhnle, 2015, 2016a,b).

### 1.2 Study Hypotheses and Objective

The guiding hypotheses contended here is that the dynamic patterns of thinking or inferences of children with ADHD elicited by a solving problem task (computer version of a popular board game known as “Battleship”) are different when compared with their typically developing controls and that there are spatiotemporal fractal measures (i.e., dynamic descriptors) that could discriminate between individuals with ADHD from those without it. Moreover it was conjectured that these measures could be used as potential biomarkers for ADHD, and serve in the future to implement a diagnostic, screening and monitoring test.

### 1.3 Working Hypotheses

i. The mean of the fractal measures from the ADHD group will be different from the typically developing control group when compared with a *MannWhitney U test* at the *p* = 0.05 significance level.

ii. Using a set of fractal measures, in a supervised classification model, aimed to identify ADHD from non-ADHD cases, the area under the receiver operating characteristics ROC curve (AUC) should be *ϑ* ≥ 0.80.

### 1.4 Objective

The main objective of this clinical trial is aimed to answer the following phase I and phase II questions, which are directly linked to the working hypotheses.

Phase I question: Do patients with ADHD have different results, for the same set of fractal measures, from non-ADHD individuals?

Phase II question: Among patients in whom the presence or absence of ADHD is clinically known, does a classifier based on a set of fractal measures, distinguish those with and without ADHD?

To answer these questions the following steps were taken:

1. The dynamic patterns of thinking of patients with ADHD and their controls were recorded using a novel and straightforward clinical testing paradigm.

2. Geometrical representations of the dynamic patterns of thinking of patients with ADHD and their control were build.

3. Fractal measures from the former geometrical representations were determined.

4. The mean values of the fractal measures results among patients known to have ADHD and patients known not to have it were compared.

5. Several supervised classification models were built using the above fractal measures as features.

6. The diagnostic accuracy of each of the classification models were estimated by evaluating their sensitivity, specificity and the (ROC) curve area (AUC) in a “training” and in an independent “testing”, validating dataset.

## 2 Methods

### 2.1 Ethics Procedures

The research protocols and procedures for this research were fully reviewed and approved by the Central Health and Disability Ethics Committee (HDEC) from New Zealand. This decision was made through the “HDEC-Full Review Pathway”. Maori consultation for supporting this research was undertaken following the principles of the Treaty of Waitangi, acknowledging inter-cultural differences to assure that partnership, participation and protection were fulfilled.

Children and parents participating in this study, were presented with a detailed description and aims of this research. Informed assent was obtained from children and informed consent from parent(s) or guardian(s). A clause concerning consent to publication of the results of the participant was included in both documents.

#### Universal Trial Number for the Study (UTN)

U1111-1146-3085 2

#### Australian New Zealand Clinical Trials Registry

ACTRN12614000306617

### 2.2 Study Design

This study is a prospective, exploratory, phase I and phase II diagnostic accuracy study with a case-control sampling design (Knottnerus and Buntinx, 2008; Larner, 2015). In clinical research this kind of trials are meant to be the first line of testing of a new technology, before to advance to more costly and clinically oriented designs. The main objective of these studies is to determine if a new technology has any diagnostic value, it is the first step during the validation of potential biomarkers in translational research. Moreover, the rationale for choosing the participants sample is that if the new technology cannot distinguish the “sickest” from the “wellest” individuals, then there is no need to continue with the new technology assessment (Zhou et al., 2011). For the purposes of this trial and according to the Diagnostic and Statistical Manual of Mental Disorders (DSM-5) (2013), the “combined presentation” of ADHD could be considered as the “sickest” presentation of the syndrome. Complementary to this distinction, typically developing children could be labelled as the “wellest” individuals.

Despite that the treatment and analysis of the data gathered for this research is strongly guided by a machine learning approach to predictive modelling (Breiman, 2001; Shmueli, 2010; Yarkoni and Westfall, 2017; Rosenberg et al., 2018), this research consider statistic control(Miller and Chapman, 2001) for the potential confounding factors, age and gender including them as input features (co-variates) in the predictive models. This choice is supported by empirical facts that shows that including them as features eventually could produce better classification models, and no extra effort is required to collect this data during the clinical interview (Rao et al., 2015, 2017). Measures of intelligence like IQ scores were not included, due to the fact that strong logical, statistical and methodological arguments have been raised against this practise in studies of neurodevelopmental disorders, particularly in ADHD (Dennis et al., 2009; Bayard et al., 2018). Besides, the marginal predictive power that IQ scores could bring to the classification models does not justify their inclusion, due to the clinical cost in terms of time and the extra work required to collect this measures. From a translational point of view, this additional cost could become a strong deterrent to the future clinical inception of this kind of diagnostic procedures.

Following the “Standards for Reporting of Diagnostic Accuracy Studies” (STARD) 2015 initiative (Cohen et al., 2016), it is important to report that the data collection for this research was planned before that the index test and the reference standard were performed.

### 2.3 Participants

The participants for this clinical trial were recruited from the Nelson Marlborough community at the Paediatrics Department from the Nelson Marlborough District Health Board in New Zealand. The recruitment process was carried out from May 2014 through September 2017. A sample of 18 children (5–13 years old) with ADHD combined presentation, along with their typically developing controls were recruited in a consecutive series, (N=36). The age class was defined considering neurobiological findings of age-related structural changes in the brain that suggest substantial developmental changes before the age of 12 and after (Sowell et al., 2004, 2003; Giedd et al., 1999). The total sample was split in two independent groups, i.e. “training” (the data coming from this group was exclusively used to build, tune and validate the predictive model in a machine-learning scenario) and “testing” (the data from this group was exclusively used to test the performance of the model). From the total sample of (36), 26 subjects were randomly assigned to the “training” group (13 cases and 13 controls) and 10 subjects (5 cases and 5 controls) to the “testing” group. The eligibility criteria for the participants was determined as follows:

#### Inclusion criteria for cases

- A full DSM-5™ diagnosis of ADHD combined presentation with associated impairment in at least two settings.
- Conners 3^rd^rdedition™ Rating Scale hyperactivity rating greater than two standard deviations above age-and sexspecific means for the parents and the teaching version.
- ADHD patients totally naive of any psychoactive drug nor received any psychoactive therapy.

#### Exclusion criteria for cases

- Any evidence of medical or neurological disorders, or any other Axis I psychiatric disorder.

#### Inclusion criteria for controls

- No DSM-5-™ diagnosis of ADHD any presentation.
- No Conners 3^rd^edition™ Rating scale hyperactivity greater than two standard deviation above age- and sex-specific means in the parents or the teacher version.

#### Exclusion criteria for controls

- DSM-5-™ diagnosis of ADHD any presentation.
- Conners 3^rd^ edition™ Rating scale hyperactivity greater than two standard deviation above age- and sex-specific means in the parents and the teacher version.
- Any evidence of medical or neurological disorders, or any other Axis I psychiatric disorder.

#### Recruitment process

The recruitment process, was performed by the paediatricians (N=8), from the Paediatrics Outpatient Clinics from the Nelson Hospital. Currently, children are referred by the local community general practitioners to the paediatrics department with behaviour concerns or with the specific question of whether ADHD is present or not. If from the referral ADHD appears to be a possibility, the paediatricians reviewing the referrals will send out a set of standard questionnaires, these include the Conners 3^rd^ edition™ for parents and teachers. When the child was seen for the first time by the paediatricians, the Conners 3^rd^ edition™ questionnaires for parents and teachers were available to be used as part of the standard diagnostic interview, which included: the full review of the clinical history plus a physical and neurological examination. If the diagnosis of ADHD was confirmed, strategies to help with ADHD were offered and usually medication was started. The study was explained to families (parents or guardians and children) at this clinic appointment, after the diagnosis of ADHD combined presentation had been made. If they wished to be involved, they were asked not to start medication immediately, and written information was given about the study and the computer game (index test). The families were allowed time to read this material before committing to the study.

The paediatricians contacted the family to book a time for a second interview in a maximum of two weeks following diagnosis, but preferably within a few days to not delay the medication too long. If after reading the information and having a chance to discuss this with the researchers at the end of the first or in the second interview, the family did not wish to participate, standard care was started. If they consented, then the child was invited to play the game in this session. Following the game, standard care was initiated with the child starting their medication if this was planned.

Regarding the recruitment of control group, internal advertisement posters and leaflets were printed and distributed between staff members from the Nelson Health Board (NHB) inviting their children Age (5-13) to participate in the study as controls. This group went throughout the same clinical assessment performed to cases group, including Conners 3^rd^ edition™ for parents and teachers.

#### Data handling and record keeping

Participant’s confidentiality was maintained at all times using a coding system and the data was password encrypted with GnuPG 1.4.13 (GNU Privacy guard tool for secure communication and data storage), archived in TAR format and stored in a USB hard-drive. The USB hard-drive was securely stored at the research site; backup copies were stored in a password-protected computer data server. The database was updated and its integrity checked every time new data flowed in.

### 2.4 Test Methods

#### Reference standard

The reference standard used to establish the presence of the target condition in this study was a combination of a clinical interview and behavioural lists, i.e., clinical interview, which included: the full review of the clinical history plus a physical and neurological examination, Conners 3^rd^ edition™ for parents and teachers (Kollins et al., 2011) and DSM-V-™ diagnosis of ADHD combined presentation. The reference standard was chosen according to the current clinical protocols used in the Paediatrics Outpatient Clinics from the Nelson hospital for the diagnosis of ADHD. The rationale for choosing this set of procedures rests on the efficiency for clinical and research purposes and its comparable validity to other alternative diagnostic procedures. All the paediatricians assessing the reference standard were blinded to the index test results.

#### Index test

In the second interview, after receiving a comprehensive description of the study, written informed assent/consent was obtained from the children and parents. The child was presented with an inference problem-solving task, i.e., the Battleship Game (see Fig 2.1).

**Fig. 2.1.**
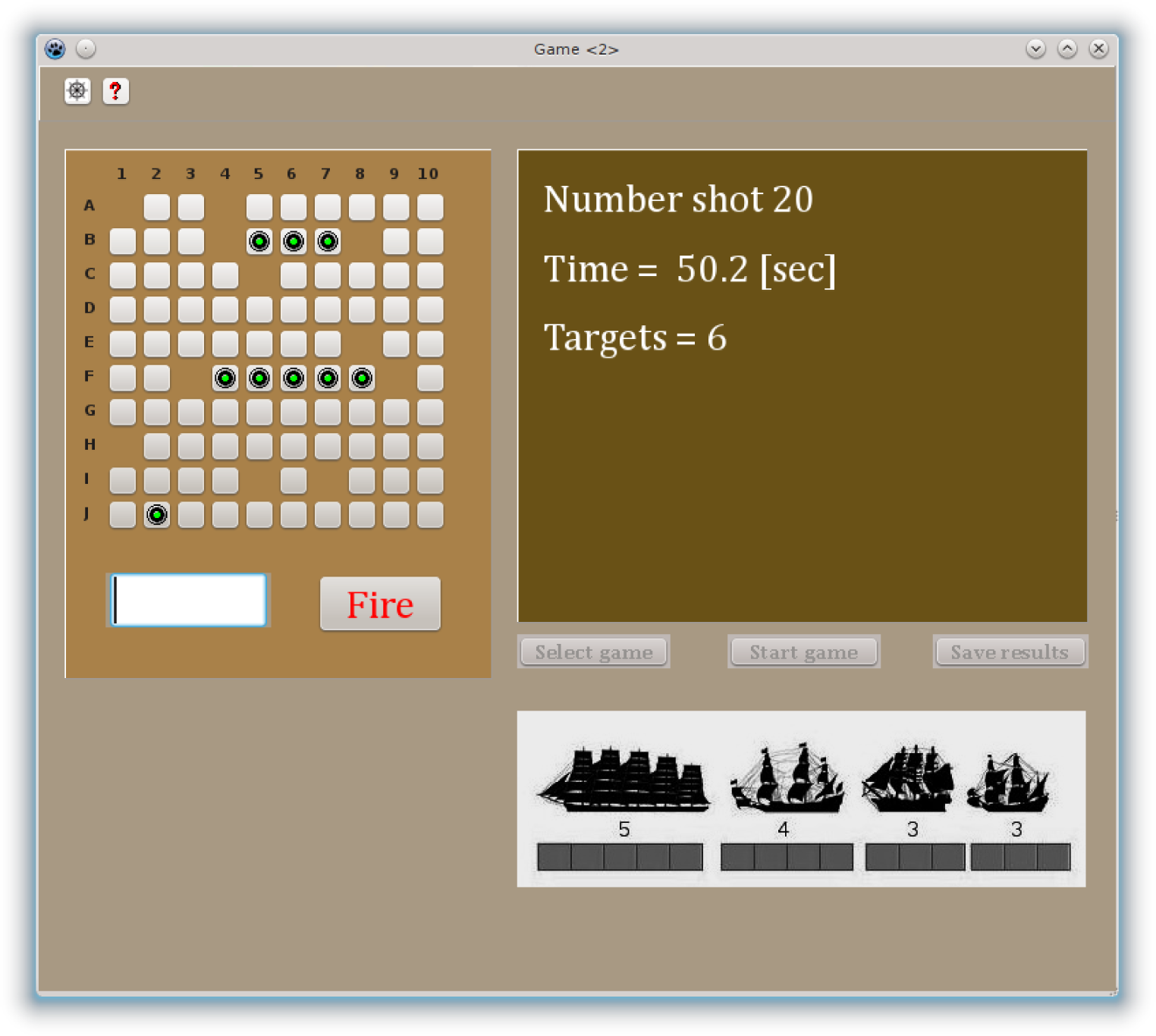
Computer Screenshot of the Battleship Game. The goal of this game was to sink all the pirates’s battleships that were hidden on the board using the least possible shots, irrespective of time used. The participants were instructed to use the pointer of the computer (mouse) to select the square that he/she wanted to fire at and then to press the Fire button. If no part of any of the ships was underneath the square selected, an empty square displayed and the sound of a water splash was heard. If any portion of a ship was hit, a green button on the grid was displayed and the sound of an explosion was heard. The right part of the screen showed a window with the number of shots taken, the time passed in seconds, and the amount of targets squares left. The bottom right part of the screen shows the silhouette of the hidden ships and the number of squared occupied by each one of them.

Battleship is a popular worldwide guessing game for two players. The original objective of the game is to find and sink all of the other player’s hidden ships before they sink all of your ships. This requires the players to devise their own battleship positions while guessing that of the other player’s. Our version of the game has been designed to be played by only one player at a time. In our case, the objective of the game is to find and sink four ships of different lengths (hidden in a board divided by a 10 × 10 grid) using the least possible shots, regardless of the time taken.

The computer version of the battleship game was developed for this research using Lazarus IDE 1.02 for the open source Pascal compiler, Free Pascal (FPC) 2.6.0, and executed under Linux Slackware 14.0. The full task includes eight individual games, each one defined by a standard template with the position of the ships (see Fig 2.2).

**Fig. 2.2.**
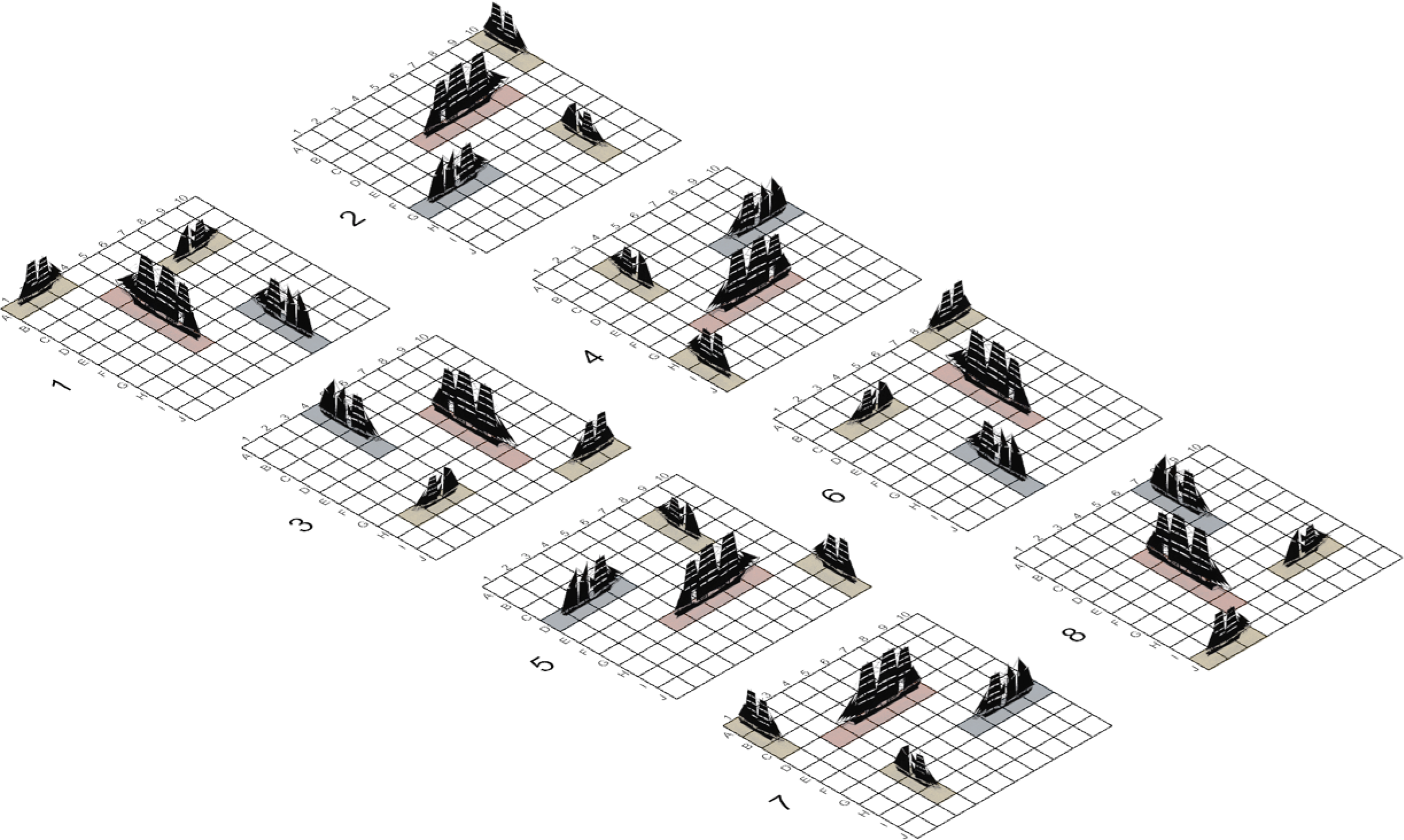
Templates with the positions of the ships on the board. The different positions were obtained according to the following procedure, the template 1 was sequentially rotated a clockwise quarter turn obtaining position 2, 3, and 4. The next four positions were obtained as specular images of positions 1, 2, 3 and 4 respectively.

Following the APA Standards for educational and psychological testing (2014) and prior to beginning each trial the interviewer initialised the computer game session with a case identification number and the current date for each participant. Afterwards the interviewer read a standardised version of the instructions (see appendix) and demonstrated to the children how to play the game in the computer.

The child was requested to find the position of four hidden ships by clicking the mouse pointer, giving a best guess regarding the position of the ships in the board. The tasks were preceded by a short practise trial with some general examples. During the task completion, the child received visual and sound feedback (in the computer screen and its speakers) about the number of shots already performed, time passed, and their ongoing performance (amount of targets left). At the end of each game the child was asked if he or she would like to play another game. The total testing duration was approximately 20-40 min; including one break after the fourth game. After finishing each game, a CSV file was created containing three columns of data: the number of shots performed, the time between shots and the (x,y) coordinated of each shot.

A Fortran code developed for this research was used to calculate Euclidean and kinematic measures from the raw CSV tabular data (distance between [x,y] coordinates, cumulative time and distance, instant speed and average final speed by shot. A second Fortran code was used to process the raw CSV tabular data to represent the time intervals between shots in a (x,y) bi-dimensional data array. The procedure consisted in interpolate a certain number of points in proportion to the amount of time used by the subject to get to the next (x,y) coordinates from the precedent shot. This procedure was iterated for the whole series of shots performed in each game (see Fig 2.3).

**Fig. 2.3.**
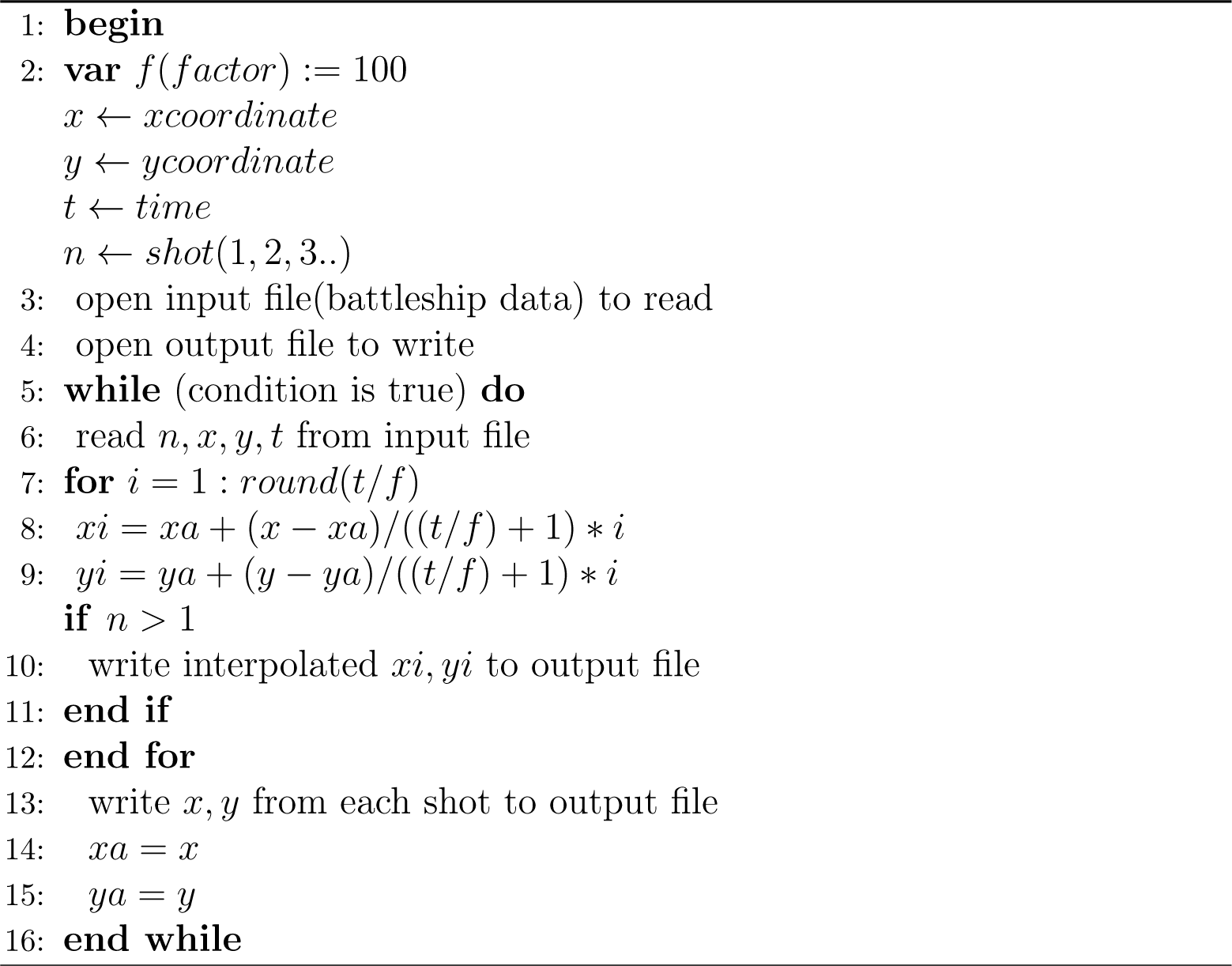
Pseudo-code for data interpolation used to build image representation of the inferential dynamics.

The final result is a new CSV file containing two columns of interpolated (x,y) coordinates.

Using this former interpolated CSV file as input, an image representing the inferential dynamics was built by plotting the (x,y) coordinates. For this purpose, a Gnuplot (Version 4.4 patchlevel 3) script was used to create a standard graphic file in PNG format. The resulting PNG file is simply a binary image of (2048 ™ 2048) pixels, without borders. This can be shown see examples in Fig 2.4) as a black background with the pixels of interest (representing the interpolated points) in white colour.

**Fig. 2.4.**
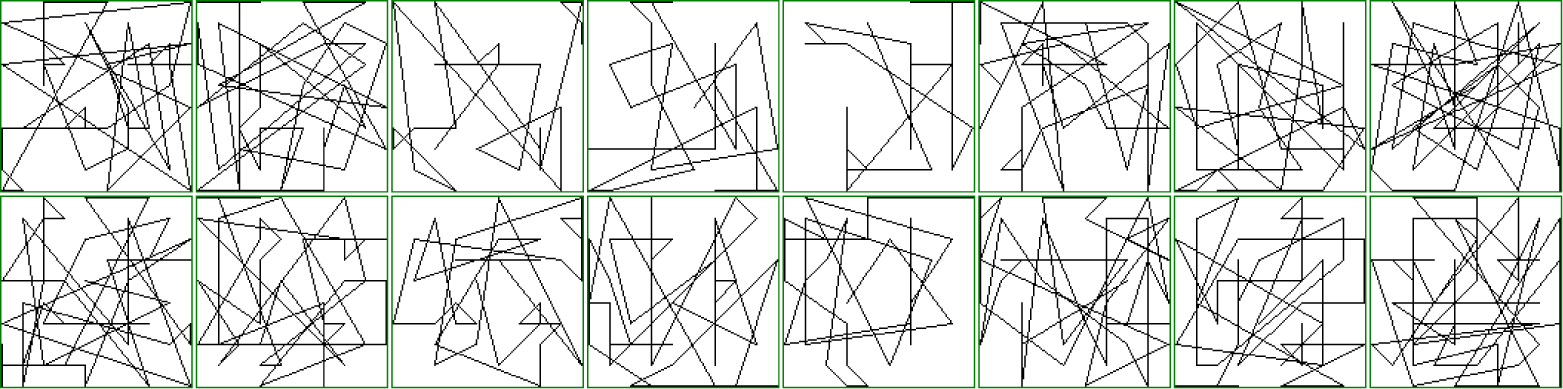
B&W color inverted example of images representing the inferential dynamics. This figure displays the 8 games played by two participants. The images from the top row correspond to subject N1 and the bottom row to subject N2. Each column represents a game ranging from the 1^st^ to 8^th^.

### 2.5 Analysis

#### Sample size calculations

The intended sample size for this research, was obtained after performing a calculation based on a 2-tailed test with type (I) error rate (α, Significance) = 0.05 and type (II) error (β, 1-power) = 0.20 and considering the conjecture that the area under the ROC curve AUC θ ϑ 0.80, [conjecture based on ADHD artificial simulated data performance (Labra-Spröhnle, 2016a)] and with a data-set ratio of normal to abnormal cases (x)=1 (Hanley and McNeil, 1982; Pepe, 2004). After this calculation, 13 participants (5–13 years) with ADHD along with 13 controls, (thus N=26); were recruited from the community. Additionally, to fulfil machine learning requirements in order to predict the unbiased error of the final results and the performance of the classification models on any new data (Williams, 2011), 5 more ADHD participant and their controls were added as a testing dataset, (N=10). The size of the testing dataset follows the commonly accepted rule of thumb employed in machine learning studies, that is to use approximately 1/3 of the size of the training dataset (Lewis, 2017). The total sample size for this research is (thus N=36).

#### Fractal Analysis and Fractal/Multiscale Measures

Fractal analysis has become a popular kind of pattern analysis that has been successfully used to investigate phenomena relevant to neuroscience (John et al., 2015). This kind of analysis is well suited to assess complex patterns across a range of different temporal and spatial scales of observation, i.e., their multiscale structure. In particular, fractal theory and its associated analytical tools have been used to describe the static and dynamics of complex spatio-temporal patterns of neural structures and cognitive functions. Notwithstanding the importance of this topic we can not provide here a full account of it. For the interested reader a good introductory text is “The Fractal Geometry of the Brain” by Leva (2016).

Briefly, fractal or multiscale measures are measures that quantifies the complexity of a given spatial or temporal pattern. Loosely speaking, they gives us a measure of the form (the geometric signature) of the pattern, allowing the identification and comparison between different forms. Based on this measure’s capabilities, we trialled the diagnostic potential of the following measures in this research:

#### (i)Multifractal Analysis

The multifractal measures used in this research were calculated from the graphic PNG file using IQM-2.01_alpha_2013_02_27 (Interactive Quantitative Morphology) (Kainz et al., 2015). These measures consisted of a Renyi spectrum of Generalised Dimension from *q=0* to *q=10*.

Multifractal analysis is a generalisation of fractal analysis and was first applied to problems of turbulence (Sreenivasan, 1991). This is a fractal method that describes complex heterogeneous spatio-temporal patterns which, unlike mono-fractals patterns, cannot be completely described with just one fractal dimension and require a full spectrum of dimensions (Seuront, 2010). This spectrum of dimensions, generally represented either as a *D*_*q*_ versus *q* plot (the moment order *q* is any number in the range -*∞* to *∞* (Feder, 1988)), or as the equivalent *f(α) α* spectrum. The dimensions *D*_*q*_ have been called the generalised dimensions, or the Renyi dimensions (Hentschel and Procaccia, 1983).

To estimate *D*_*q*_ the method of moments based on the box-counting algorithm could be used (Halsey et al., 1986). In this method an image to be analysed is covered with a grid, which divided it into *N(ϵ)* squares of side length ϵ, allowing calculation of the mass *m*_*i*_*(ϵ)* in each of them. After this step the partition function could be computed:

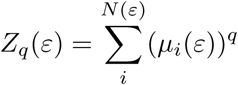

The operation is performed for different values of ϵ and *q*, within a predetermined range. The generalized dimension *D*_*q*_ is calculated as:

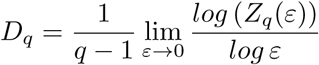

The term *D*_*q*_ is calculated as the slope of the *log(Z*_*q*_) versus *log(ϵ)*; this is done for different *q*. Using a range of *q* that includes negative and positive numbers produces a graph of *D*_*q*_ in terms of *q*, the so called spectrum of generalised dimensions or Renyi dimensions (Saravia et al., 2012).

Several algorithms have been developed to obtain the spectrum of generalised dimension or Renyi spectrum (Seuront, 2010). In this research, the raster box algorithm was used due to its simplicity and accurate performance for the estimation of the positive side of the spectrum (Mach et al., 1995; Saa et al., 2007).

The Renyi dimensions were calculated using IQM running in a Linux Slackware64, 14.0 machine using the following parameters:

1. Raster box algorithm option was selected
2. Minimum *q* = [0]
3. Maximum *q* = [10]
4. 4 # eps = [12]. This number corresponds to the 12 different side lengths ϵ of the boxes used in the raster box algorithm. The values of length ϵ in pixels are: 1, 2, 4, 8, 16, 32, 64, 128, 256, 512, 1024, 2045.
5. Regression: Start = [6]; End = [12]. This values indicate the box sizes selected for the estimation of each *D*_*q*_ (by computing the slope of the *log(Z*_*q*_) versus *log(ϵ)* in the linear regression) that is, from boxes sized 32 pixels to boxes sized 2045 pixels.

#### (ii)Lacunarity Analysis

The lacunarity measures were also calculated from the graphic PNG files, already used in the multifractal analysis. Lacunarity analysis is a fractal technique for describing patterns of temporal and spatial dispersion. This method is widely applicable to many data set used in the human and natural sciences. Although originally developed for mono-fractal objects, the method in its general form, can be used to describe multifractal patterns. (Plotnick et al., 1996). This method can be applied to data of any dimensionality and from multiples sources. It allows the description of scale dependent changes in spatial and temporal structures, by these means it can give insights of the dynamic of underlying processes. The technique has been implemented in several packages for image and signal analysis (Rosenberg Michael S. and Anderson Corey Devin, 2010; Karperien et al., 2013; Kainz et al., 2015).

The gliding-box procedure for lacunarity estimation can be describes as follows: a box of length *r* is placed at the origin of one of the images. The number of occupied sites within the box (box mass = *s*) is established. The box is displaced one space along the image and the box mass is again estimated. This process is repeated over the entire image, producing a frequency distribution of the box masses *n(s,r)*. This frequency distribution is converted into a probability distribution *Q(s,r)* by dividing by the total number of boxes *N(r)* of size *r*. Then the first and second moments of this distribution are estimated:

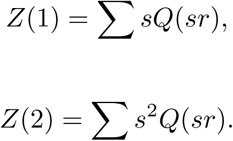

The lacunarity for this box size is now defined as:

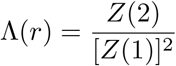

This calculation is repeated over a range of box sizes from *r* = 1 to 10. A log-log plot of the lacunarity versus the size of the gliding box is then produced (Plotnick et al., 1996).

The lacunarity measures, from Lac1 to Lac10 were also calculated with IQM using the following parameters:

1. Gliding box algorithm option was selected.
2. Maximum epsilon = [10] This correspond to the maximum size (*r*) of the gliding box side.
3. Regression: Start = [1]; End = [10]. This correspond to the range of log-log plot of the lacunarity versus the 10 different sizes of the gliding box.

#### (iii)Multiscale Straightness Index (MSSI) analysis

The MSSI measures were obtained from the CSV tabular data file using an Octave m-script, developed for this research and executed in Octave, version 3.8.1, running in a Linux Slackware-64, 14.0 machine.

The MSSI analysis is a multiscale method aimed to assess the “straightness” of a spatiotemporal trajectory multiple times, over a range of selected scales of temporal resolution (i.e., ‘granularity’) and at different location of observational ‘windows’ (Postlethwaite et al., 2013). These attributes permit the accurate description of a range of displacements of a natural or artificial moving object. The MSSI is computed by iterative sub-sampling the trajectory data at all possible temporal granularities.

Paraphrasing Postlethwaite et al (2013), let the individual location estimates of a moving object comprising trajectories be given by triplets (*x*_*j*_, *y*_*j*_, *t*_*j*_), for 0,…, *N-1* where *N* is the total number of position fixes in the track. The point (*x*_*j*_, *y*_*j*_) is the location of the moving object at time *t*_*j*_, and let *t* _0_ = 0. Also we suppose that the time interval between fixes is a constant, *s*. That is *t*_*j*+1_*t*_*j*_ = *s*, for all *j*.

The granularity, *g*, is defined as the interval at which the trajectory data is observed, and the window, *w*, is the length of time over which MSSI is computed. Generally, both *g* and *w* are integer multiples of *s*, and *w* is an integer multiple of *g*, but other values of *g* and *w* are possible by interpolating between fixes if is needed. The track spacing *s,* granularity *g* and window *w* are defined in units of time. We let:

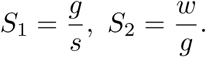

Distances between two points in a trajectory are defined by:

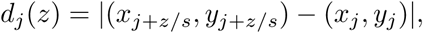

in which *z* can be either the granularity, *g*, or the window, *w*. Thus the MSSI can be defined as:

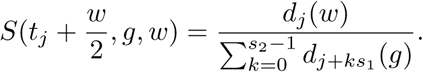

The first argument of *S* gives the time at which the MSSI is defined, the second argument is the granularity, and the third is the window’s size. The numerator of the fraction is the beeline distance between two location estimates; separated by a time interval of *w*, and the denominator is the total distance travelled between the same two locations observing the track at a time interval equal to *g*. Summing up, to compute *S,* first the track data is re-sampled at an interval equal to *g*, then the beeline distance is calculated between the position of the moving object at time *t*_*j*_, and the position at time *t*_*j*_*+ w*, where *w* is the window, *S* is the ratio of this distance to the total distance travelled by the mobile over this time window, using the re-sampled track data at a granularity *g* A value of *S* close to 1 indicates travel in a straight line. A small value of *S* indicates highly tortuous movement.

The following parameter were used in the Octave m-script to obtain the measures:

1. Granularity = [1]
2. Range of windows = [2] starting window size; [2] increment in size of the windows series; [60] final windows size.

Using this script, 30 different MSSI measures were obtained according to the different range of windows size used, i.e., (2, 4, 6, …, 60).

#### Kinematic analysis

Kinematic analysis was performed using a Fortran code developed for this research and compiled with GFortran under Linux Slackware-64, 14.0 This routine was used to calculate Euclidean and dynamics measures from the CSV tabular data file i.e., geometric distance between shots, cumulative distance, instantaneous speed and average speed per shot.

#### Mann-Whitney U test

Previous to perform means comparisons with Mann-Whitney U test, ShapiroWilk test was used to determine normality, and Levene’s test was performed to assess the equality of variances for the variables obtained from the ADHD and non-ADHD groups. The significance level for both test was set at 0.05 and their values were estimated using the R package (Kabacoff, 2015).

After summarising the metric variables: age, time, distance, number of shots and the set of multiscale measures from the eight games played by the participants in this research, Mann-Whitney U test at 0.05 significance level was carried out to compare their means; this statistic was estimated using the R package. Statistics of effect size for the Mann–Whitney test were calculated to provide standardised measures of how much difference exist between the two groups by using the Pearson’s r statistics. This measures an their confidence intervals were obtained with the “BootES” package and executed in R (Kirby and Gerlanc, 2013).

#### Classification models

The Statistical Computing Platform R, version 3.3.3 (2017-03-06) – “Another Canoe” and the “Caret” package were used to build supervised machine learning predictive models, using a multivariate set of multifractal, lacunarity, MSSI and kinematic measures. The R packages “Rattle” and “RWeka” were used to build alternative machine learning models. The performance of the predictive models and their ability to distinguish ADHD subjects from controls was determined using a probability cut-off value = 0.5.

Despite that neural networks, support vector machine and decision trees were used during the first steps in determining the best classification models, these machine learning techniques are not reviewed here, this task exceeds the limits and the scope of this paper. Moreover excellent texts exist where the interested reader can further investigate this point (i.e. Kuhn and Johnson, 2013).

**Logistic regression** Logistic regression models are very popular classifiers due to their simplicity and efficiency when the response is categorical and the main goal is solely prediction/classification, specially in a repeated measures scenario (Cleophas and Zwinderman, 2013). Models based on logistic regression are aimed at predicting the probability of an event in subjects at risk. Nonetheless, the method requires the user to identify an effective pool of predictors (feature selection) that yield its best performance. The task of producing logistic regression models was performed using the R package “caret” implemented by Kuhn (2013).

**Features selection** Using the full set of variables, the selection of predictors for the models was performed using the R package “fscaret” (Automated Feature Selection from “caret” (Szlęk, 2016)) and the models were further tuned manually using a forward selection strategy (the departing model always included variables age and sex) to get the highest values of AUC (ROC) in the training dataset using a repeated 10-fold cross-validation procedure (Kuhn and Johnson, 2013).

**Control for the confounding variables age and sex** To prevent residual confounding, due to the unknown type of association (linear or non linear) between the confounding variables (age and sex) and the outcome from the classification models, control for both confounding were performed. The adjustment for the continuous confounding variable age, was done by means of stratification in five categorical strata following empirical suggestions derived from clinical studies (Groenwold et al., 2013; Cochran, 1968). The categories of age in years ranged from 0<(a)=<8<(b)=<9<(c)=<10<(d)=<11<(e); this confounding variable was included as a co-variate factor in the predictive models. The main reason for controlling for age using stratification in categories is due to the fact that the underlying processes of cognitive development are not continuous and they can rise or decline at different ages (Werner, 1957). For the confounding variable sex, a dichotomous category was used and also included as a co-variate factor in the predictive models (Rao et al., 2015, 2017).

**Analysis of diagnostic accuracy** Receiver Operating Characteristics (ROC) analysis was used to calculate several performance measures including, accuracy, sensitivity, specificity and the ROC AUC (Swets, 2014). These tasks were performed using the “Caret” and “pROC” R packages (Kuhn, 2008; Robin et al., 2011).

**Confidence intervals and p-values for ROC analysis** Confidence intervals CI, at a significance level of 0.05 were calculated for ROC AUC, sensitivity and specificity using a bootstrapping technique implemented in the “pROC” R package (Swets, 2014). Calculations of CI and p-values at a significance level of 0.05 for the accuracy were performed using an in-house, bootstrapping R script produced for this research.

## 3 Results

### 3.1. Participants

#### Participants demographics

The participants baseline demographic and clinical characteristic are presented in Table 1. A total of 36 participants were recruited, of whom 18 were ADHD cases and 18 non-ADHD controls. The age range of the overall sample was 5-13 years (average age in months = 105.50 and standard deviation = 19.48). The total sample consisted of 14 females and 22 males. The total sample was split in 2 datasets, the training dataset (n= 26) and the testing dataset (n=10). Both datasets were balanced having the same amount of ADHD cases and controls.

**Tab. 1.**
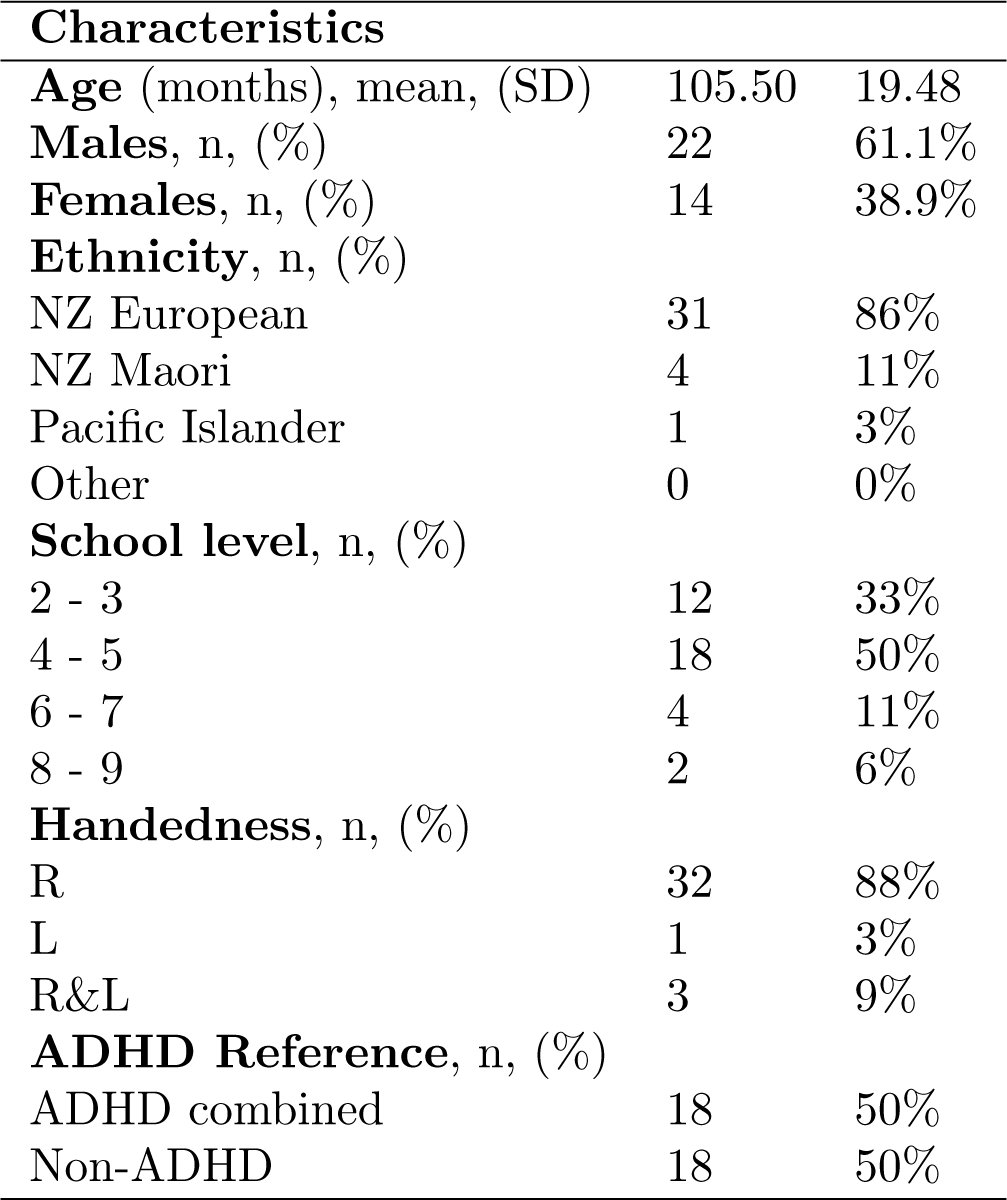
Demographic and Sample Characteristics

| **Characteristics** |  |  |
| --- | --- | --- |
| **Age** (months), mean, (SD) | 105.50 | 19.48 |
| **Males**, n, (%) | 22 | 61.1% |
| **Females**, n, (%) | 14 | 38.9% |
| **Ethnicity**, n, (%) |  |  |
| NZ European | 31 | 86% |
| NZ Maori | 4 | 11% |
| Pacific Islander | 1 | 3% |
| Other | 0 | 0% |
| **School level**, n, (%) |  |  |
| 2 - 3 | 12 | 33% |
| 4 - 5 | 18 | 50% |
| 6 - 7 | 4 | 11% |
| 8 - 9 | 2 | 6% |
| **Handedness**, n, (%) |  |  |
| R | 32 | 88% |
| L | 1 | 3% |
| R&L | 3 | 9% |
| **ADHD Reference**, n, (%) |  |  |
| ADHD combined | 18 | 50% |
| Non-ADHD | 18 | 50% |

#### Flow of participants

The flow of participants was initiated by the community general practitioner and it is thoroughly detailed in two flow diagrams contained in figs 3.1 and 3.2. A supplementary STARD diagram provides additional information in Fig 3.3. As the STARD diagramm shows, 24 participants drop out of the study during the performance of the index test. From these, 20 participant belonging to the control group and 4 to the cases group. The cause for this drop out in the control group happens because the participant were performing the test during lunch time break, with a restrictive amount of time to fulfill the whole task composed by 8 games. In the cases group, two drop outs were due to malfunction of the recording software, and two by the negative of the child to finish the full task.

**Fig. 3.1.**
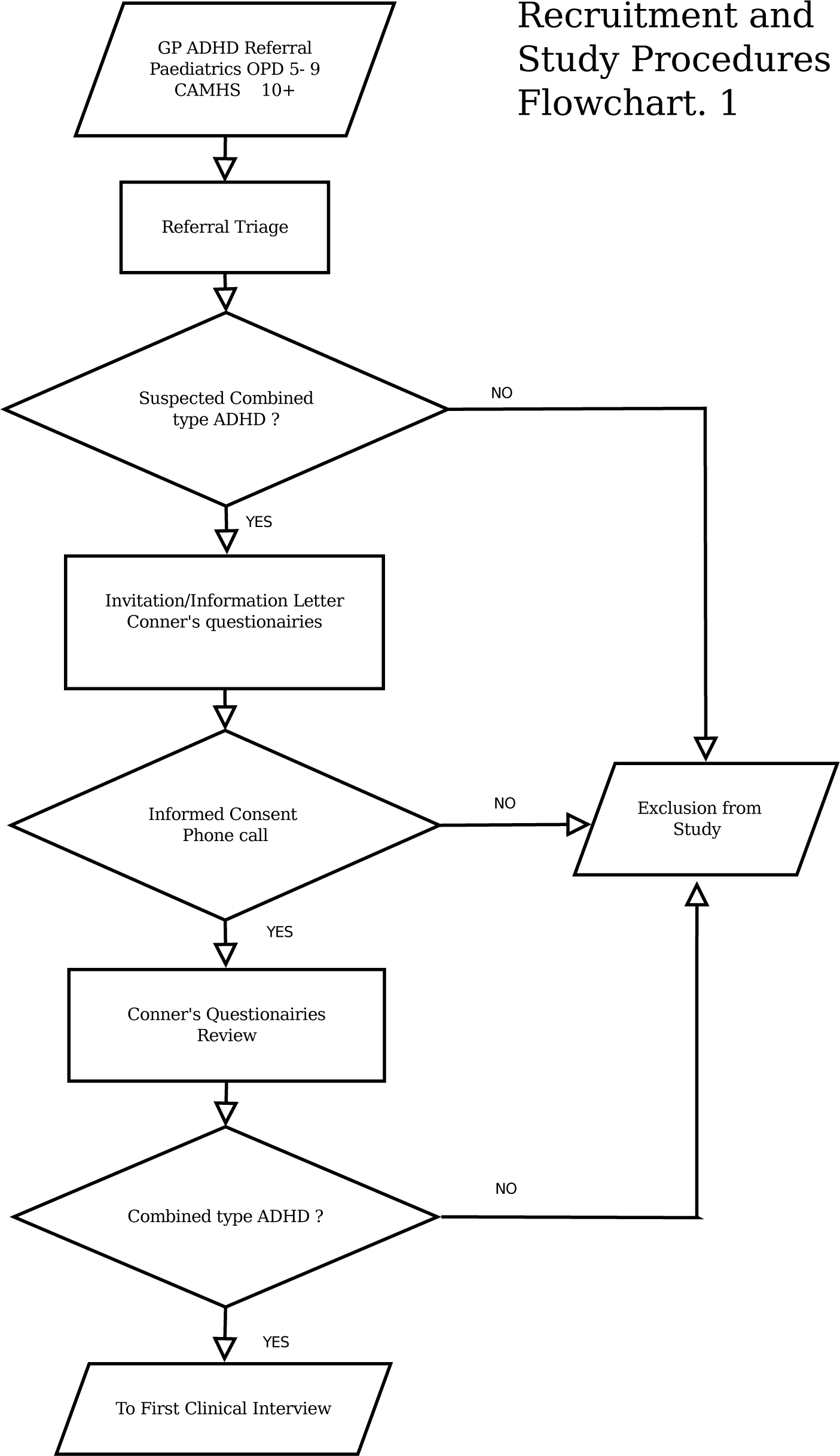
Study procedures flowchart N1

**Fig. 3.2.**
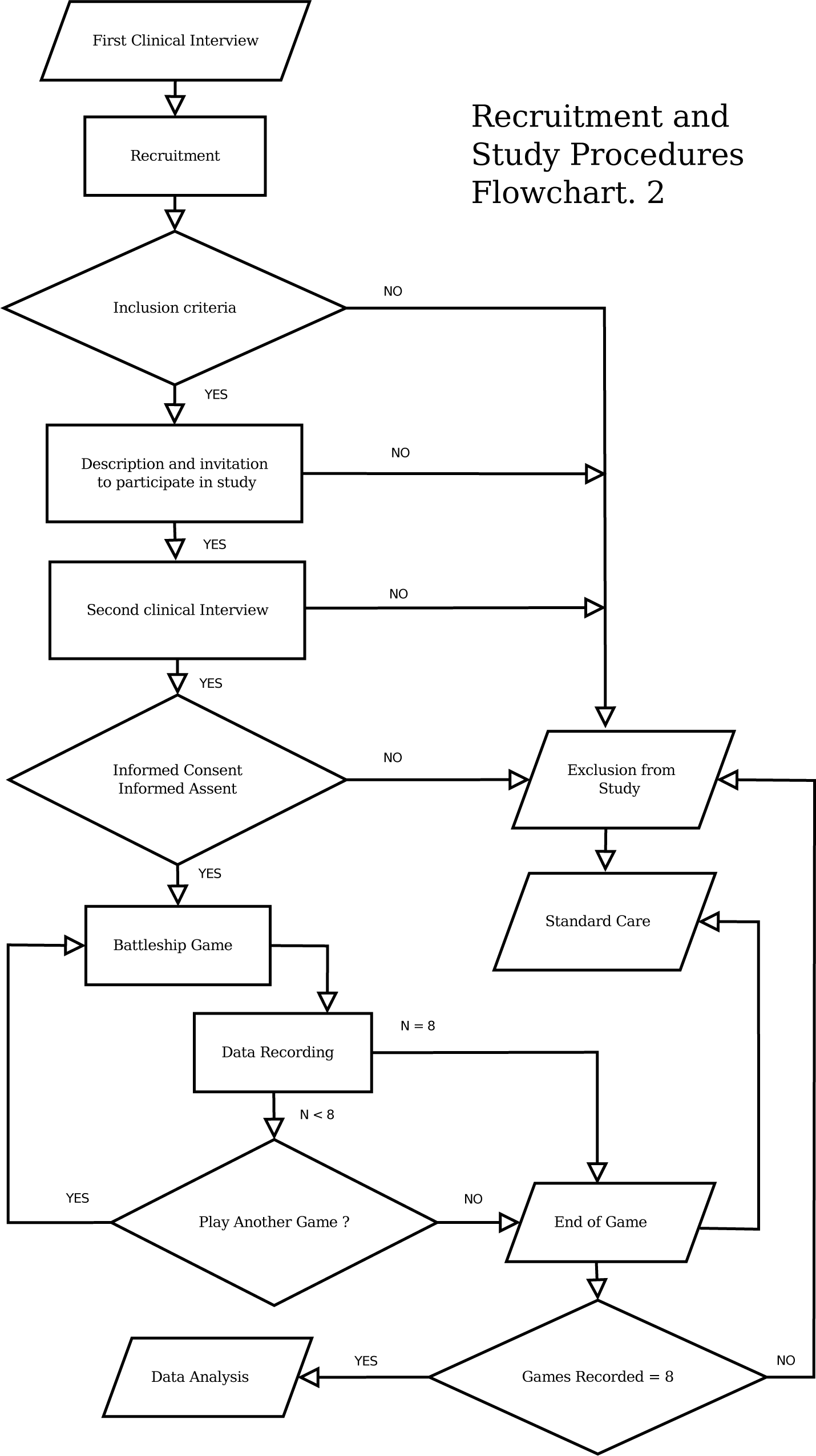
Study procedures flowchart N2

**Fig. 3.3.**
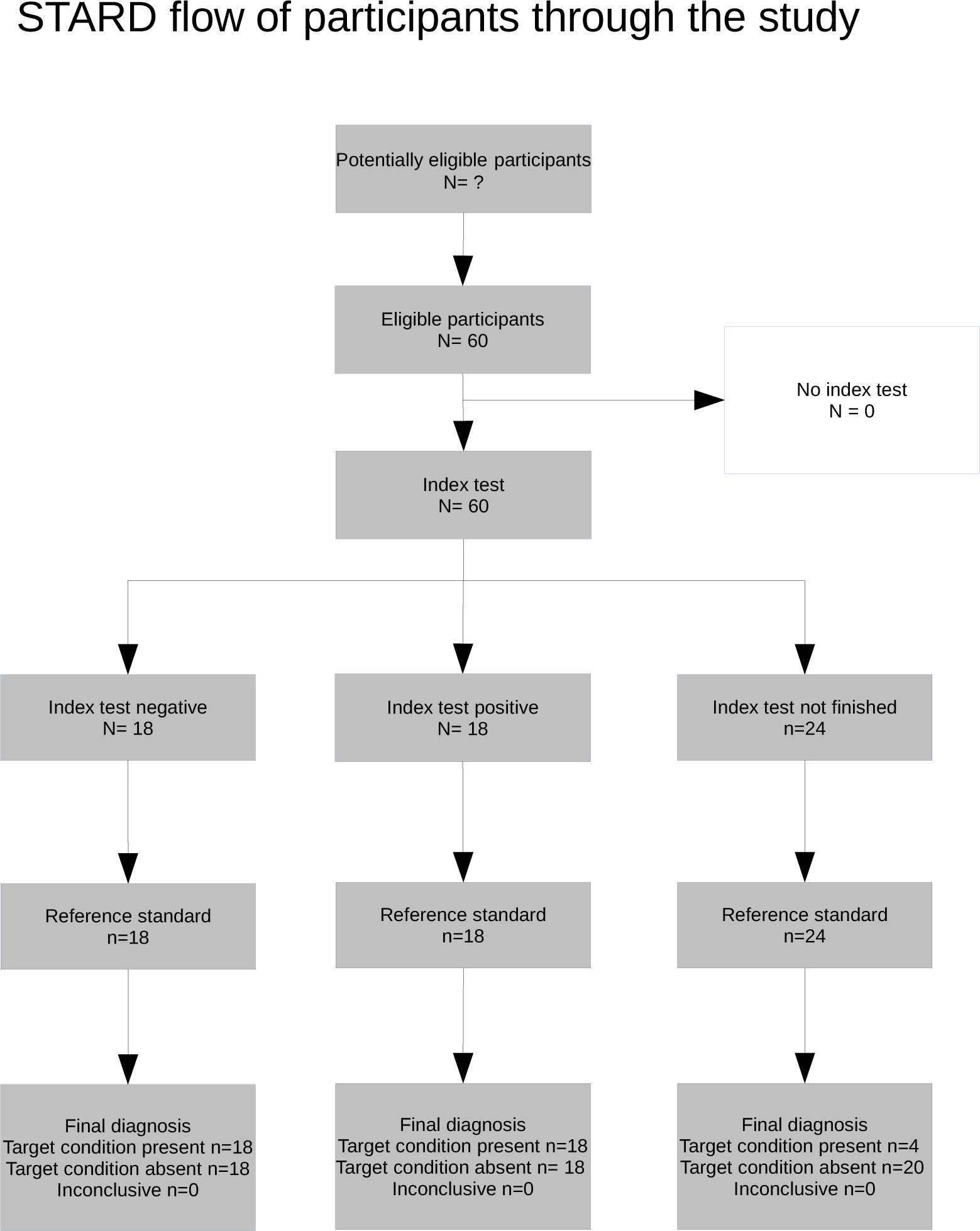
STARD Flow of participants through the study

### 3.2. Test Results

No adverse events from performing the index test or the reference standard happened during the trial. The results are presented following the primary objective of this clinical diagnostic research and that aimed to answer the phase I and II questions contained in the next subsections.

### 3.3 Reference Standard

#### A. Cases group

The application of the reference standard to the cases group produced very homogeneous results in all the instances, confirming the presence of the ADHD combined presentation according to the full review of the clinical history, the physical and neurological examination, the Conners 3^rd^ edition™ for parents and teachers and the DSM-V-™.

#### B. Control group

The application of the reference standard to the control group confirmed the absence of ADHD in any of its clinical presentations, but produced more heterogeneous results than the cases group. In one case, hyperactivity was detected by the Conners 3^rd^ edition™ for teachers but not for parents. Nevertheless the clinical history, the physical and neurological examination and DSM-V-™ ruled out the presence of ADHD in this group.

### 3.4 Index Test

#### Results for the Phase I Question

The visual inspection of the histograms with the distribution of age, time, distance and the multiscale measures shows conspicuous differences for ADHD and non-ADHD individuals (see Figure 3.4).

**Fig. 3.4.**
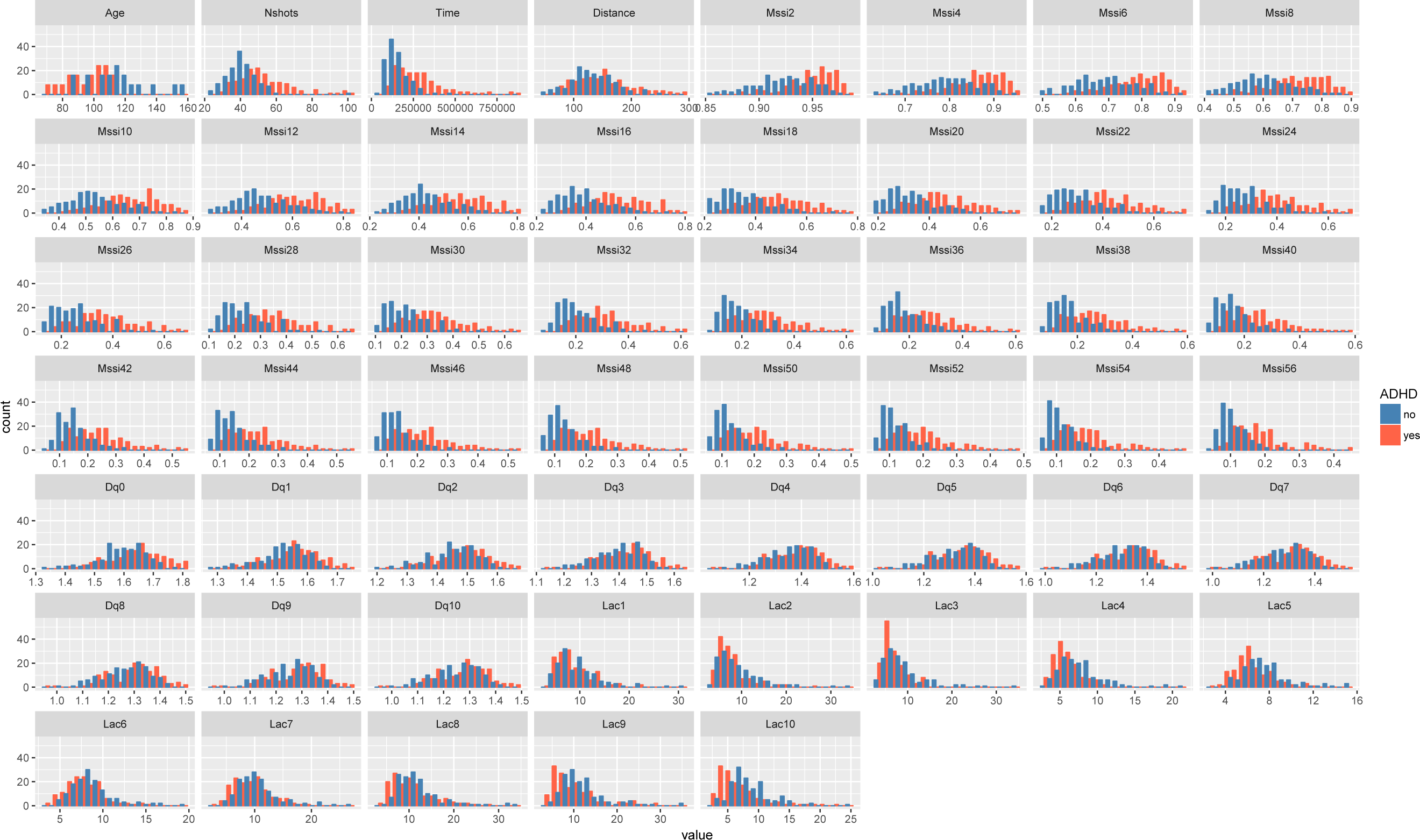
Answering the Phase I question. Histograms with the distribution of age, number of shots (Nshots), time, distance and multiscale measures [multifractal (Dq), lacunarity (Lac) and Multiscale Straightness Index (Mssi)] for ADHD (red) and Control group (blue).

From the histograms is also apparent a non-normal distribution of the variables which is compatible with the results of the Shapiro-Wilk test (see Tables 2 and 3).

**Tab. 2.**
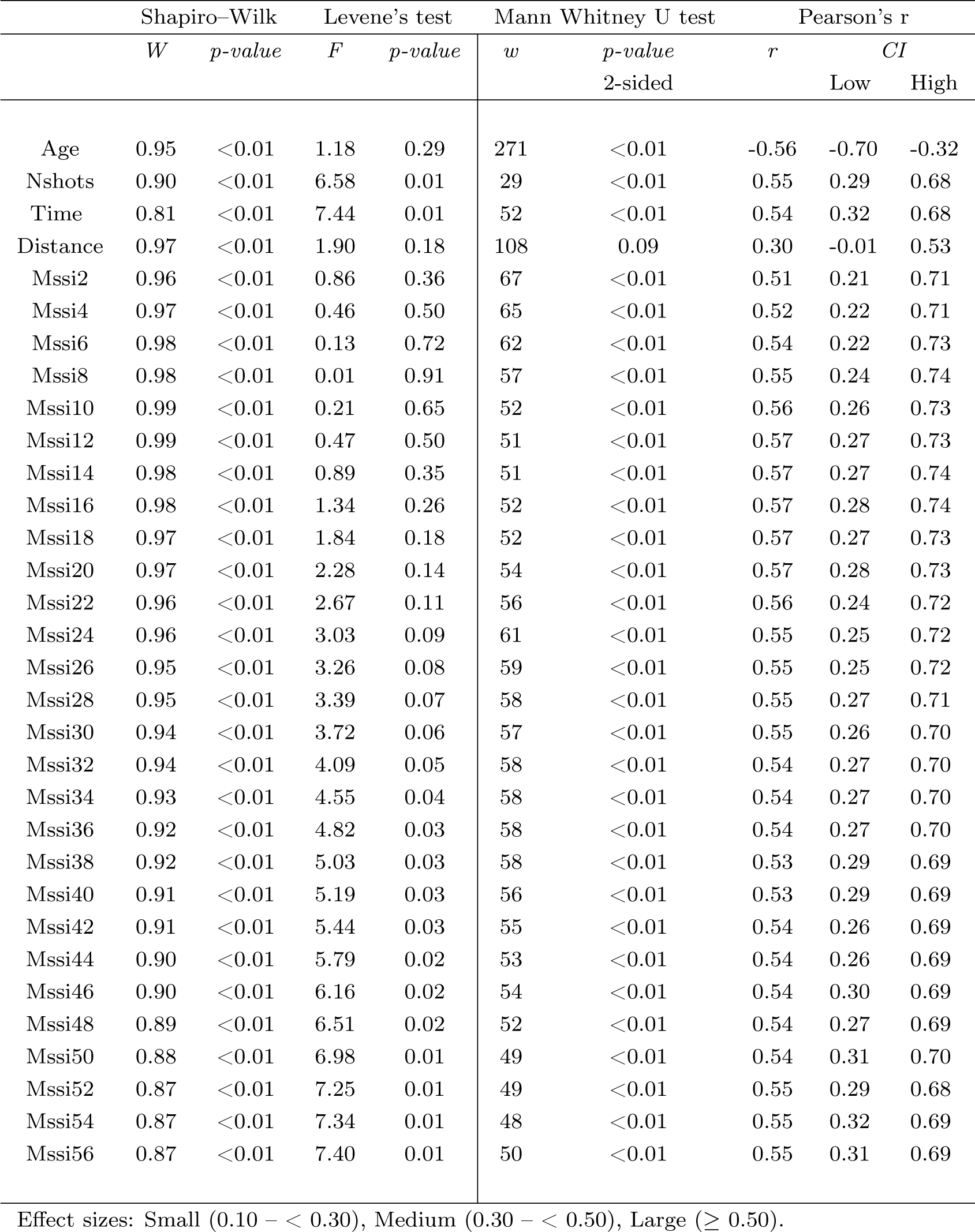
Answering the Phase I question. Comparing the means of age (months), number of shots, time (millisec), distance and multiscale measures (multiscale straightness index (MSSI)) for the ADHD and Control group with the Mann-Whitney U test and the effect size statistics Pearson’s r. Shapiro-Wilk test (W), and Levene’s test values (F) and their respective p-values are reported to assess normality and the equality of variances between ADHD and control groups.

**Tab. 3.**
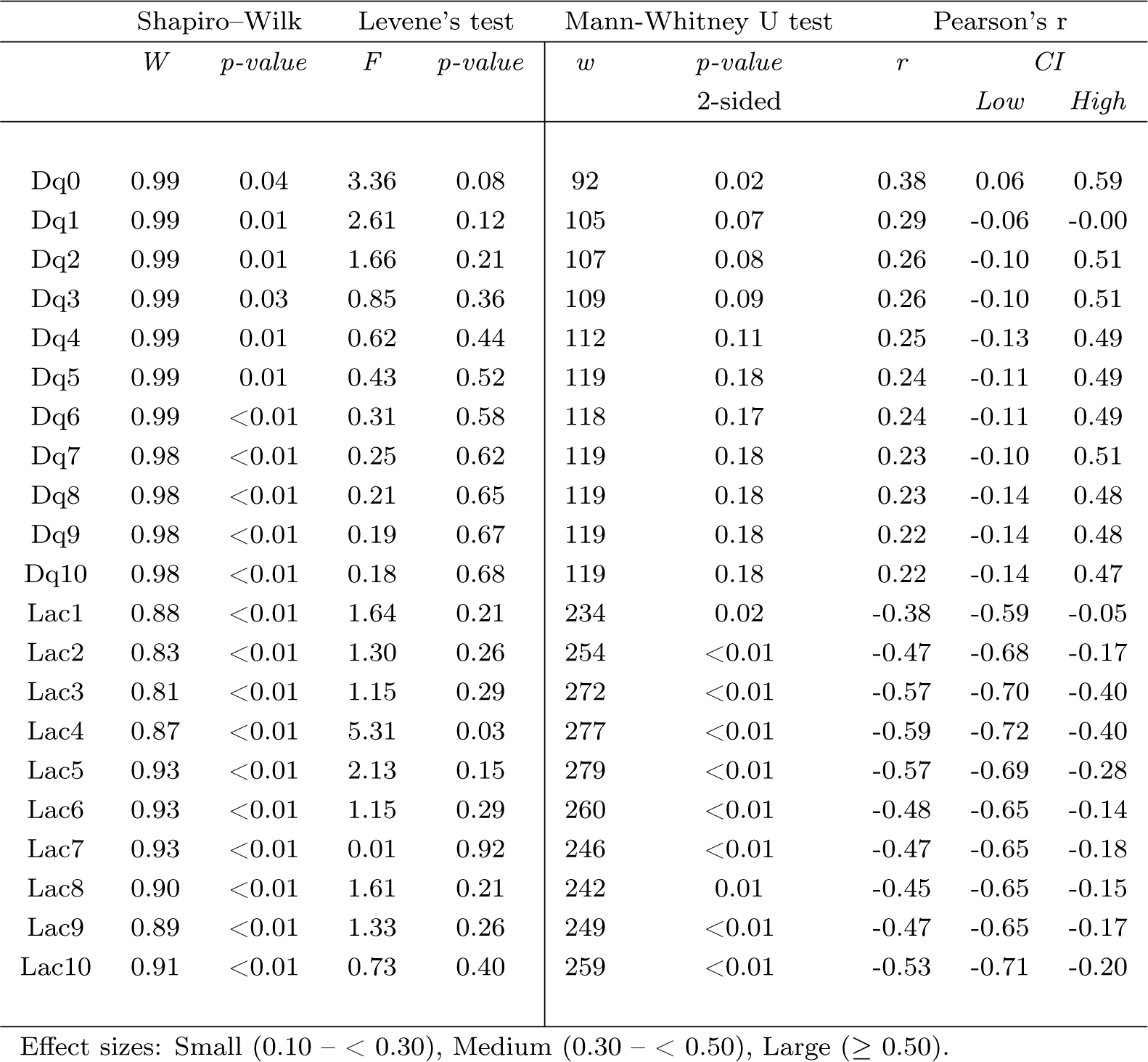
Answering the Phase I question. Comparing the means of Multiscale Measures (Generalised Rényi dimensions’ spectrum and Lacunarity spectrum) for the ADHD and Control group with the Mann-Whitney U test and the effect size statistics (r). Shapiro-Wilk test (W) and Levene’s test values (F) and their p-values are reported to assess normality and the equality of variances between ADHD and control groups.

Results from the Mann-Whitney U test, at significance level 0.05, shows that the variables, age (months) (Time) in millisec and number of shots (Nshots) display significant differences across ADHD diagnosis. The variable distance (Dis) display non-significant differences. Most of the means of the fractal measures from the ADHD and the control group are different. The (MSSI) measures present the more conspicuous differences across ADHD diagnosis. The lacunarity measures (Lac) also show significant differences across ADHD categories. In contrast with this findings, most of the multifractal measures (Dq) display non-significant differences, except (Dq0). These differences are manifested with diverse strength as is shows by the effect sizes estimated with the Pearson correlation coefficient, r. From these measures it can be observed:

1. The “Large” (r *ϑ* 0.5) and significant effect size for the variables: (Age), (NShots), (Time), (Lac3), (Lac4), (Lac5), (Lac10), and all the MSSI estimates.

2. A “Medium” significant effect size (r = 0.3 – < 0.5) for the variables, (Lac1), (Lac2), (Lac6), (Lac7), (Lac8), (Lac9) and (Dq0). The variable distance (Dis) display a Medium but non-significant effect size.

3. A “Small” non-significant effect size (r = 0.1 – < 0.3) for the variables: (Dq1 to Dq10).

#### Results for the Phase II Question

Several supervised machine learning models, based on a set of fractal measures used as predictors, were built to classify the subjects, including: logistic regression, random forest, support vector machines and neural networks. However, despite an improved classification accuracy being achieved by all the machine learning models, only results for the linear logistic regression model are reported in this section. This decision was taken due to the limited number of samples available in the training dataset. The models not reported were excluded because they are prone to over-fit the training data and give overly optimistic estimates of the classification performance (Kuhn and Johnson, 2013). Additionally, these models are difficult to interpret because of their black-box nature (Kuhn and Johnson, 2013).

The logistic regression model was built in the training dataset using 11 predictors: (number of shots [Ns]/time [time], and generalised Renyi dimensions [Dq0], [Dq1], [Dq2]; MSSI measures: [mssi42], [mssi44], [mssi56]; lacunarity: [Lac1], [Lac2], [Lac3], [Lac10]), and 2 confounding variables: (age and sex).

The diagnostic accuracy, sensitivity, specificity and the (ROC) curve area (AUC) for the logistic regression model are presented in the figure 3.5 and tables 4 and 5. The logistic regression classification (risk) probabilities (for the training and testing datasets) and its mean by individuals are presented in tables 6 and 7.

**Fig. 3.5.**
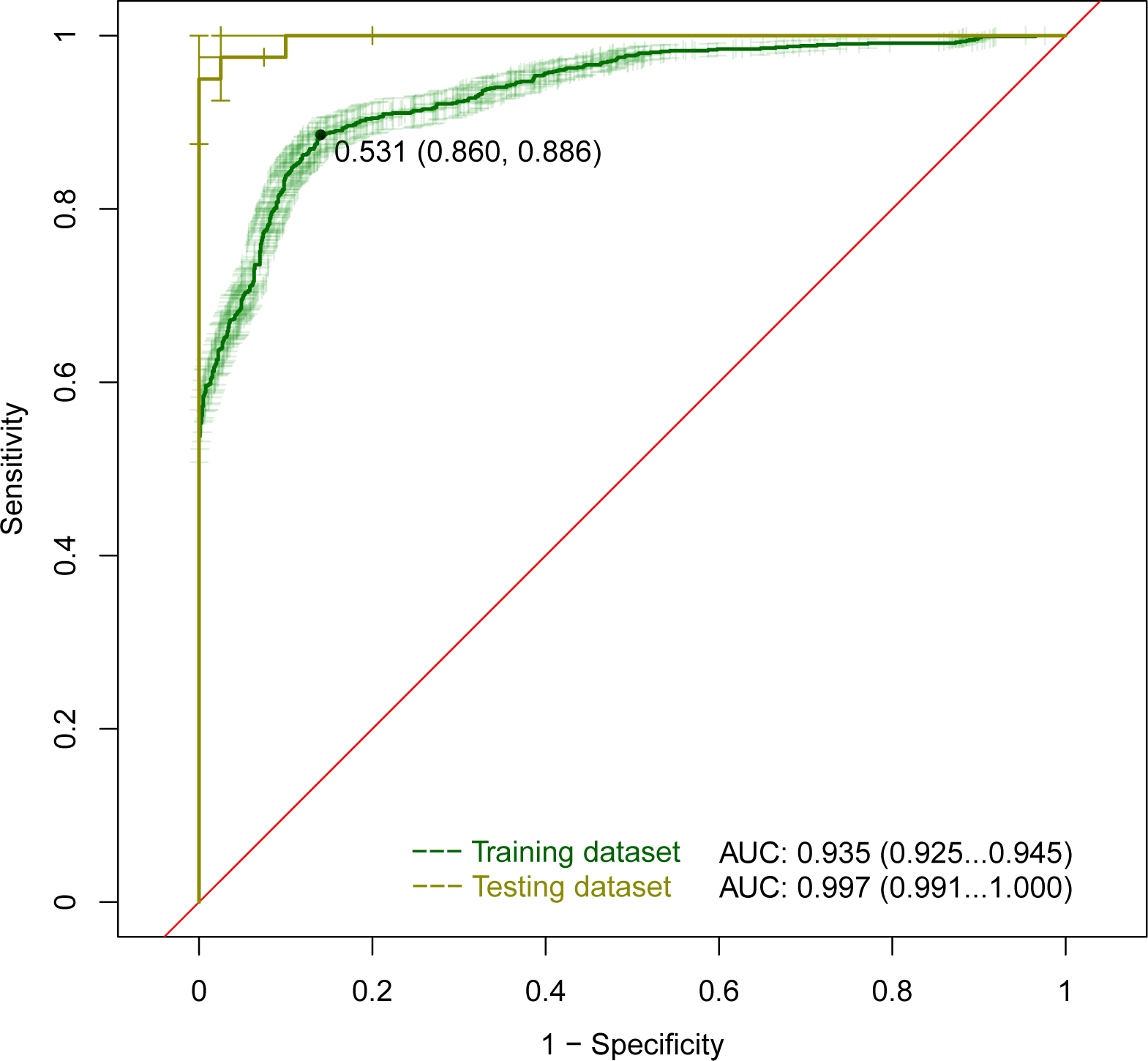
(ROC) curve area (AUC) for the naive logistic regression classifier running in the training dataset [95% CI for ROC curve (shaded in green) and optimum cut-off value included in the plot]. ROC AUC and its 95% CI for the testing dataset are also reported in this plot.

**Tab. 4.** Answering the Phase II question in the **training dataset**. Among patients in whom the presence or absence of ADHD is clinically known, does a classifier based on a set of fractal measures, distinguish those with and without ADHD? The results display the performance, at the level of separates games (instances), of a classifier using a naive logistic regression. A resampling scheme with a repeated 10-fold crossvalidation was used to produce an appropriate estimate of the model performance in the training dataset. The 95% confidence intervals for the accuracy values were obtained using a resampling R-script with 1000 bootstrap samples.

| Confusion Matrix and Statistics for Training Dataset |  |  |
| --- | --- | --- |
| Prediction | Reference |  |
|  | ADHD | No-ADHD |
| ADHD | 901 | 133 |
| No-ADHD | 139 | 907 |

| Classification characteristics and their 95% confidence intervals |  |  |
| --- | --- | --- |
|  | Lower | Upper |
| Accuracy = 0.8692, p-value <2e-16 | 0.846 | 0.951 |
| ROC (AUC) = 0.935 | 0.925 | 0.945 |
| Sensitivity = 86.63% | 85.44% | 89.54% |
| Specificity = 87.21% | 85.95% | 89.98% |
| Positive Predictive Value = 87.14% | 86.15% | 89.67% |
| Negative Predictive Value = 86.71% | 85.78% | 89.31% |

**Tab. 5.** Answering the Phase II question in a independent **testing dataset**. The results display the performance, at the level of separates games (instances), of the former tuned naive logistic regression classifier running in the testing dataset. The 95% confidence intervals for the accuracy values were obtained using a resampling R-script with 1000 bootstrap samples.

| Confusion Matrix and Statistics for Testing Dataset |  |  |
| --- | --- | --- |
| Prediction | Reference |  |
|  | ADHD | No-ADHD |
| ADHD | 40 | 4 |
| No-ADHD | 0 | 36 |

| Classification characteristics and their 95% confidence intervals |  |  |
| --- | --- | --- |
|  | Lower | Upper |
| Accuracy = 0.95, p-value = 0.003 | 0.875 | 1.000 |
| ROC (AUC) = 0.997 | 0.991 | 1.000 |
| Sensitivity = 100.00% | 90.97% | 100.00% |
| Specificity = 90.00% | 76.34% | 97.21% |
| Positive Predictive Value = 90.91% | 79.37% | 96.11% |
| Negative Predictive Value = 100% |  |  |

**Tab. 6.**
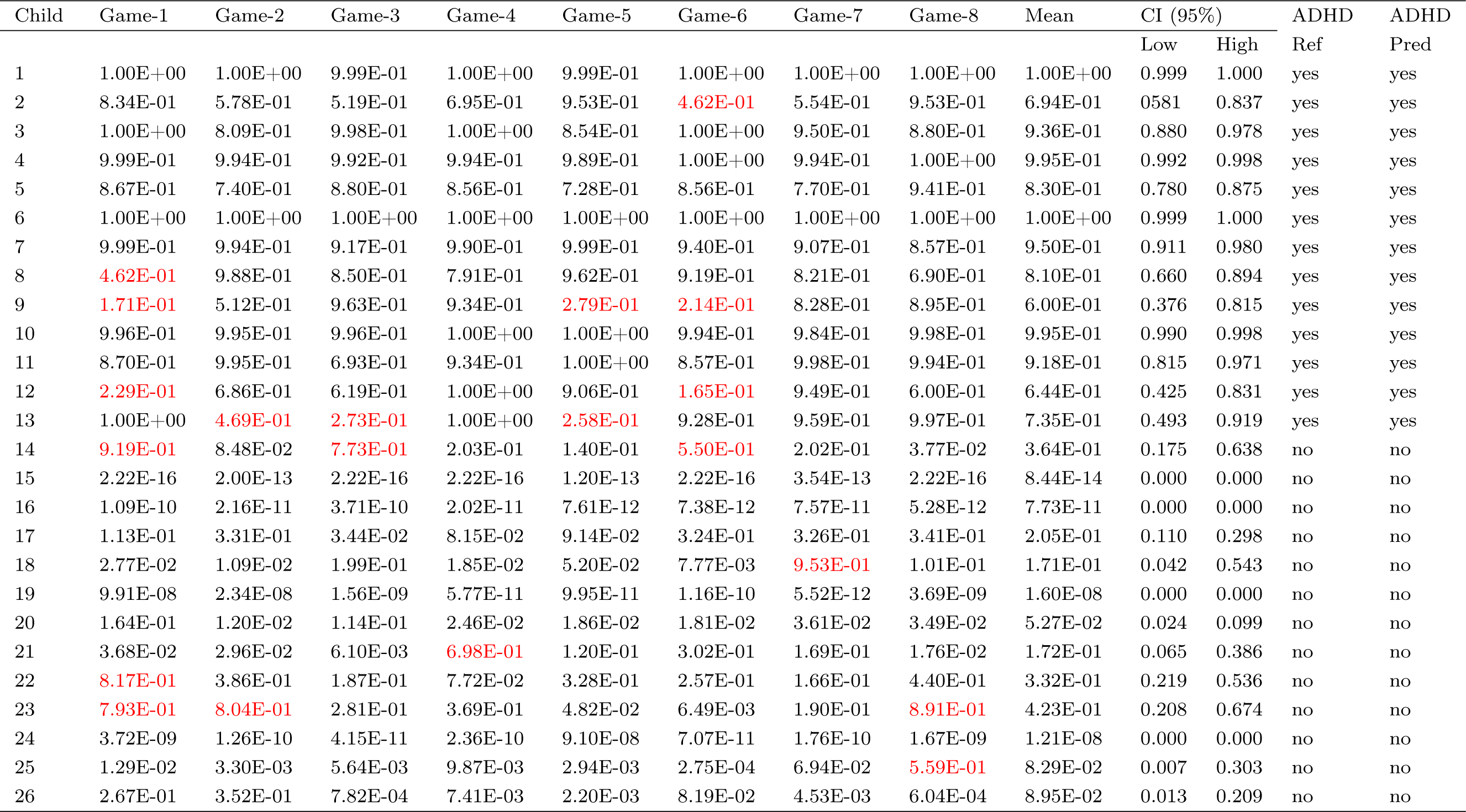
Logistic regression classification risk probabilities from the training dataset: individual games classification probabilities and mean with their 95% CI per participant across their games played. Predictions for the positive class (ADHD presence) were determined using a probability cut-off value of the mean = 0.5. Highlighted in red are the individual game predictions (Pred) not matching the reference standard (Ref).

| Child | Game-1 | Game-2 | Game-3 | Game-4 | Game-5 | Game-6 | Game-7 | Game-8 | Mean | CI (95%) |  | ADHD Ref | ADHD Pred |
| --- | --- | --- | --- | --- | --- | --- | --- | --- | --- | --- | --- | --- | --- |
|  |  |  |  |  |  |  |  |  |  | Low | High |  |  |
| 1 | 1.00E+00 | 1.00E+00 | 9.99E-01 | 1.00E+00 | 9.99E-01 | 1.00E+00 | 1.00E+00 | 1.00E+00 | 1.00E+00 | 0.999 | 1.000 | yes | yes |
| 2 | 8.34E-01 | 5.78E-01 | 5.19E-01 | 6.95E-01 | 9.53E-01 | **4.62E-01** | 5.54E-01 | 9.53E-01 | 6.94E-01 | 0.581 | 0.837 | yes | yes |
| 3 | 1.00E+00 | 8.09E-01 | 9.98E-01 | 1.00E+00 | 8.54E-01 | 1.00E+00 | 9.50E-01 | 8.80E-01 | 9.36E-01 | 0.880 | 0.978 | yes | yes |
| 4 | 9.99E-01 | 9.94E-01 | 9.92E-01 | 9.94E-01 | 9.89E-01 | 1.00E+00 | 9.94E-01 | 1.00E+00 | 9.95E-01 | 0.992 | 0.998 | yes | yes |
| 5 | 8.67E-01 | 7.40E-01 | 8.80E-01 | 8.56E-01 | 7.28E-01 | 8.56E-01 | 7.70E-01 | 9.41E-01 | 8.30E-01 | 0.780 | 0.875 | yes | yes |
| 6 | 1.00E+00 | 1.00E+00 | 1.00E+00 | 1.00E+00 | 1.00E+00 | 1.00E+00 | 1.00E+00 | 1.00E+00 | 1.00E+00 | 0.999 | 1.000 | yes | yes |
| 7 | 9.99E-01 | 9.94E-01 | 9.17E-01 | 9.90E-01 | 9.99E-01 | 9.40E-01 | 9.07E-01 | 8.57E-01 | 9.50E-01 | 0.911 | 0.980 | yes | yes |
| 8 | **4.62E-01** | 9.88E-01 | 8.50E-01 | 7.91E-01 | 9.62E-01 | 9.19E-01 | 8.21E-01 | 6.90E-01 | 8.10E-01 | 0.660 | 0.894 | yes | yes |
| 9 | **1.71E-01** | 5.12E-01 | 9.63E-01 | 9.34E-01 | **2.79E-01** | **2.14E-01** | 8.28E-01 | 8.95E-01 | 6.00E-01 | 0.376 | 0.815 | yes | yes |
| 10 | 9.96E-01 | 9.95E-01 | 9.96E-01 | 1.00E+00 | 1.00E+00 | 9.94E-01 | 9.84E-01 | 9.98E-01 | 9.95E-01 | 0.990 | 0.998 | yes | yes |
| 11 | 8.70E-01 | 9.95E-01 | 6.93E-01 | 9.34E-01 | 1.00E+00 | 8.57E-01 | 9.98E-01 | 9.94E-01 | 9.18E-01 | 0.815 | 0.971 | yes | yes |
| 12 | **2.29E-01** | 6.86E-01 | 6.19E-01 | 1.00E+00 | 9.06E-01 | **1.65E-01** | 9.49E-01 | 6.00E-01 | 6.44E-01 | 0.425 | 0.831 | yes | yes |
| 13 | 1.00E+00 | **4.69E-01** | **2.73E-01** | 1.00E+00 | **2.58E-01** | 9.28E-01 | 9.59E-01 | 9.97E-01 | 7.35E-01 | 0.493 | 0.919 | yes | yes |
| 14 | **9.19E-01** | 8.48E-02 | **7.73E-01** | 2.03E-01 | 1.40E-01 | **5.50E-01** | 2.02E-01 | 3.77E-02 | 3.64E-01 | 0.175 | 0.638 | no | no |
| 15 | 2.22E-16 | 2.00E-13 | 2.22E-16 | 2.22E-16 | 1.20E-13 | 2.22E-16 | 3.54E-13 | 2.22E-16 | 8.44E-14 | 0.000 | 0.000 | no | no |
| 16 | 1.09E-10 | 2.16E-11 | 3.71E-10 | 2.02E-11 | 7.61E-12 | 7.38E-12 | 7.57E-11 | 5.28E-12 | 7.73E-11 | 0.000 | 0.000 | no | no |
| 17 | 1.13E-01 | 3.31E-01 | 3.44E-02 | 8.15E-02 | 9.14E-02 | 3.24E-01 | 3.26E-01 | 3.41E-01 | 2.05E-01 | 0.110 | 0.298 | no | no |
| 18 | 2.77E-02 | 1.09E-02 | 1.99E-01 | 1.85E-02 | 5.20E-02 | 7.77E-03 | **9.53E-01** | 1.01E-01 | 1.71E-01 | 0.042 | 0.543 | no | no |
| 19 | 9.91E-08 | 2.34E-08 | 1.56E-09 | 5.77E-11 | 9.95E-11 | 1.16E-10 | 5.52E-12 | 3.69E-09 | 1.60E-08 | 0.000 | 0.000 | no | no |
| 20 | 1.64E-01 | 1.20E-02 | 1.14E-01 | 2.46E-02 | 1.86E-02 | 1.81E-02 | 3.61E-02 | 3.49E-02 | 5.27E-02 | 0.024 | 0.099 | no | no |
| 21 | 3.68E-02 | 2.96E-02 | 6.10E-03 | **6.98E-01** | 1.20E-01 | 3.02E-01 | 1.69E-01 | 1.76E-02 | 1.72E-01 | 0.065 | 0.386 | no | no |
| 22 | **8.17E-01** | 3.86E-01 | 1.87E-01 | 7.72E-02 | 3.28E-01 | 2.57E-01 | 1.66E-01 | 4.40E-01 | 3.32E-01 | 0.219 | 0.536 | no | no |
| 23 | **7.93E-01** | **8.04E-01** | 2.81E-01 | 3.69E-01 | 4.82E-02 | 6.49E-03 | 1.90E-01 | **8.91E-01** | 4.23E-01 | 0.208 | 0.674 | no | no |
| 24 | 3.72E-09 | 1.26E-10 | 4.15E-11 | 2.36E-10 | 9.10E-08 | 7.07E-11 | 1.76E-10 | 1.67E-09 | 1.21E-08 | 0.000 | 0.000 | no | no |
| 25 | 1.29E-02 | 3.30E-03 | 5.64E-03 | 9.87E-03 | 2.94E-03 | 2.75E-04 | 6.94E-02 | **5.59E-01** | 8.29E-02 | 0.007 | 0.303 | no | no |
| 26 | 2.67E-01 | 3.52E-01 | 7.82E-04 | 7.41E-03 | 2.20E-03 | 8.19E-02 | 4.53E-03 | 6.04E-04 | 8.95E-02 | 0.013 | 0.209 | no | no |

**Tab. 7.** Logistic regression classification risk probabilities from the testing dataset. Individual games classification probabilities and mean with their 95% CI per participant across their games played. Predictions for the positive class (ADHD presence) were determined using a probability cut-off value of the mean = 0.5. Highlighted in red are the individual game predictions (Pred) not matching the reference standard (Ref).

| Child | Game-1 | Game-2 | Game-3 | Game-4 | Game-5 | Game-6 | Game-7 | Game-8 | Mean | CI (95%) |  | ADHD Ref | ADHD Pred |
| --- | --- | --- | --- | --- | --- | --- | --- | --- | --- | --- | --- | --- | --- |
|  |  |  |  |  |  |  |  |  |  | Low | High |  |  |
| 27 | 9.73E-01 | 9.92E-01 | 9.96E-01 | 9.97E-01 | 8.17E-01 | 9.98E-01 | 1.00E+00 | 9.81E-01 | 9.69E-01 | 0.881 | 0.993 | yes | yes |
| 28 | 8.91E-01 | 9.99E-01 | 1.00E+00 | 1.00E+00 | 1.00E+00 | 9.98E-01 | 1.00E+00 | 9.27E-01 | 9.77E-01 | 0.936 | 0.999 | yes | yes |
| 29 | 1.00E+00 | 1.00E+00 | 1.00E+00 | 1.00E+00 | 1.00E+00 | 9.59E-01 | 9.54E-01 | 9.98E-01 | 9.89E-01 | 0.972 | 1.000 | yes | yes |
| 30 | 1.00E+00 | 1.00E+00 | 9.71E-01 | 9.99E-01 | 9.82E-01 | 9.83E-01 | 9.83E-01 | 9.98E-01 | 9.90E-01 | 0.982 | 0.996 | yes | yes |
| 31 | 1.00E+00 | 1.00E+00 | 1.00E+00 | 1.00E+00 | 1.00E+00 | 1.00E+00 | 1.00E+00 | 1.00E+00 | 1.00E+00 | 1.000 | 1.000 | yes | yes |
| 32 | 2.87E-01 | 3.72E-02 | 7.14E-03 | 5.49E-02 | **7.25E-01** | 7.42E-03 | 1.62E-03 | 2.84E-04 | 1.40E-01 | 0.031 | 0.457 | no | no |
| 33 | 4.82E-02 | 4.40E-02 | 2.33E-01 | 8.80E-03 | 1.73E-02 | 5.67E-02 | 5.50E-04 | 7.10E-02 | 6.00E-02 | 0.028 | 0.141 | no | no |
| 34 | 9.87E-03 | 1.12E-04 | 2.48E-05 | 7.05E-06 | 1.52E-05 | 3.91E-06 | 6.05E-05 | 1.81E-05 | 1.26E-03 | 0.000 | 0.006 | no | no |
| 35 | 1.31E-07 | 1.08E-08 | 2.12E-08 | 6.00E-09 | 5.25E-09 | 2.20E-07 | 2.00E-09 | 1.52E-07 | 6.86E-08 | 0.000 | 0.000 | no | no |
| 36 | 3.23E-01 | **7.10E-01** | 2.73E-01 | **8.90E-01** | **9.54E-01** | 3.47E-01 | 1.22E-02 | 2.00E-01 | 4.64E-01 | 0.255 | 0.686 | no | no |

## 4 Discussion

To rightly situate the results of this research in the context of other existing objective methods developed to identify and classify ADHD requires distinguishing two dimensions to make comparisons; the qualitative, which point to the means (the methods, tools and their related constructs) by which the diagnostics were achieved, and the quantitative, which is referred to the measures of diagnostic accuracy.

In regards to the qualitative dimension, the comparisons were assisted by a review of meta-analyses of the neurocognitive profile of ADHD (Pievsky and McGrath, 2017), and by a systematic review of the physiological substrates of executive functioning (Munro et al., 2017). From the first review, twelve cognitive domains could be identified, these domains corresponds to the main constructs abstracted from the diverse tools and methods employed to assess ADHD. The domains identified were: decision-making; fluency; intelligence/achievement; memory; planning/organisation; reaction time; reaction time variability; response inhibition; selective attention; set shifting; vigilance, and working memory. The second review presented and analysed works intended to functionally map the former domains onto different brain structures. These two reviews are exemplar in presenting the main tenets of the mainstream cognitivist paradigm that drives current clinical and neuroscience research devoted to investigate ADHD and EF. In this paradigm, organised behaviour is conceived as underpinned by a modular psychological organisation in faculties and its functioning is operated by computational mechanism with a more or less stable localisation in different brain structures (Fodor, 1983; Carruthers, 2006). For what follows it is worth commenting that the concept of EF was born under the aegis of the Soviet dialectical-materialism (cultural-historical paradigm) and migrated from there, but in a ill-defined manner, to the Anglo-Saxon cognitivist paradigm (Labra-Spröhnle, 2016b). It was Shallice (1982), Duncan (1986), Welsh & Pennington (1988), whom in a series of very influential papers, mainly contribute to this conceptual shift. In those papers, they candidly but uncritically suggested that the EF concept, taken from Luria (1980) can be radically interpreted using an information-processing model and a machine problem solving framework (Newell and Simon, 1971). In this regard, particularly influential was the work of Welsh & Pennington whom using terminology from cognitive science, equated the EF concept to Posner’s notion (1986) of a “limited-capacity central processing system”, and also to other functions ascribed to the frontal lobes, e.g., cognitive control (Welsh and Pennington, 1988). In this manner the stage was set for the coming years of the EF conceptual controversies (more than thirty definitions have been produce since then) and the vibrant, diverse but puzzling EF research performed by the mainstream cognitive scientist. Under the cognitivist paradigm, the notion of EF was disconnected from its original sources i.e., (Anokhin and Bernstein’s functional system theory, Vygotsky and Luria’s cultural-historical approach and Filimonov’s principle of “graded and pluripotential localization of functions”), losing its integrative character and transplanted to a modular and computational view of cognition (Fodor, 1983; Carruthers, 2006). In this framework, EF was fragmented in diverse separate mental functions (faculties), underpinned by distinct, more or less localised, brain networks in which one is the seat of a central control (Uttal, 2003; Perler, 2015). This conceptual shift favoured multiple, *ad-hoc*, arbitrary “extension” to the concept of EF, with poor or null operational character without reaching consensus (Barkley, 2012). As a consequence the methodological aspect and the experimental expression of EF models were downgraded in their validity and utility (Labra-Spröhnle, 2016b). Contributing to the aggravation of the same situation, another paradoxical condition happened during cognitive revolution and despite its “revolutionary” conceptual shift, the practitioners of the new paradigm kept attached to the same old style and tools of experimental testing, i.e., the “open-loop causal model” of behaviour and mental functioning (Marken, 1988, 2009; Marken and Mansell, 2013).

For an informed reader it comes as no surprise that most of the objective methods found in the current literature, aimed to help in the ADHD diagnosis and EF assessment, and that were reviewed for this research, are based or inspired in the aforementioned cognitivist paradigm enacted by “open-loop causal models” oriented to get “performance” measures (Griffin et al., 2015; Hoskyn et al., 2017). In contrast, and this is the main difference in the qualitative dimension of comparison, the theoretical constructs and the method presented in this research are based on a different paradigm.

In general, in this alternative paradigm, the variety of human behaviour repertoire and mental functioning are conceptualised as goal-directed and self-controlled activities (Piaget, 1974; Powers, 2005). Moreover, the methodological expression of this condition could be accounted in a “close-loop experimental paradigm” (Marken, 2009; Marken and Mansell, 2013) aimed to get “morphological” measures (Labra-Spröhnle, 2015), and without resorting to a faculties ^1^ or modular framework. Instead, behaviour and mental activities are conceived as integrative dynamic phenomena, organised in “functional systems” (Vygotsky, 1965; Anokhin, 1974; Luria, 1980; Labra-Spröhnle, 2016a). Functional systems are materially open (but with a cyclical, close-loop functioning), self-organizing systems, composed of synchronised, distributed, neural, bodily structures and elements from the medium in which the agent acts purposefully. Besides, it is considered that mental activity is operated by non-representational (causal) and representational (implications) means (Piaget, 2006; Pezzulo, 2008; Piaget et al., 2013), and that methodologically, this parallel activity can be expressed in a relational framework (Piaget, 1963).

In particular, the theoretical views regarding EF follows the definition provided by Labra-Spröhnle (2016a), i.e., “executive functions are any of “those specific mechanisms of the functional system which provide for the universal physiological architecture of the behavioral act”. For the purposes of this discussion, this definition should be understood in the following way:

1. This definition postulate that “those specific mechanism” in a “functional system” correspond to inferential processes.

2. It emphasises the ubiquitous character (i.e., the universal presence) of the inferential activity at any level of biological organisation.

3. It acknowledges the “cyclic” or “close-loop” structure of the inferential activity that is present in any “functional system”.

4. The inferential activity (both at sensory-motor and representative level) is deemed central (as the engine) in the functioning of any behaviour and cognition and constitute their “executive” aspect.

5. Human EF is formed and chronologically modulated by biological developmental and socio-historical conditions.

6. The emergence and localisation of EF is assumed as a synchronised activity distributed hierarchically across many different spatiotemporal scales from the brain-body-environment interaction.

7. EF disorders could be manifested as deviations from the spatio-temporal dynamical structure (morphology) of the inferential processes of typically developing individuals.

Furthermore, this view acknowledges that the causal and implications aspect of mental activities shares a complex spatiotemporal patterns, manifested at different scales (Labra-Spröhnle, 2015, 2016a,b).

Based on these former tenets and the aforementioned definition of EF, the method (and the experimental paradigm) presented here is the first one devised to objectively and metrically asses the regulative (control) character of human inferential dynamics using a closed-loop task, and also it is the first of its kind used to assess EF disorders such as ADHD.

The testing strategy implemented in the method is in line with and supported by new evidence confirming that inferential processes is a feature of mental activity greatly affected in children with ADHD (Brunamonti et al., 2017). This methodology is also convergent with new complementary theoretical views, that shows that EF dynamics could be adequately modelled by spatiotemporal “interaction-dominant” processes (Ihlen and Vereijken, 2010; Anastas et al., 2014). Moreover, the increasing attention to dynamics measures such as reaction time variability in ADHD during EF task, is also an illustrative example of another convergent trends in ADHD research (Kofler et al., 2013, 2018)

Notwithstanding, an important epistemological caveat must be placed here to avoid misunderstandings, the *“ratio cognoscendi”* shall not be confused with the *“ratio essendi”* of the phenomena described and analysed by using multiscal-ing techniques, including those advocated in this work (Piaget, 2006). It is important to bear in mind what Labra-Spröhnle (2017) and Gigerenzer (2007) described about the complex epistemological dialectic between tools and theories and their explanatory limitations due to their mutual constructive entanglement. In regards to the quantitative dimension, and acknowledging that direct comparisons between tests may be the optimal procedure; only indirect, cursory comparisons were possible at this stage. The selected comparing metrics between tests were their accuracy, sensitive and specificity. The global task of making indirect comparison in the quantitative dimension, was assisted by a recent systematic review produced to evaluate “The clinical utility of the continuous performance test and objective measures of activity for diagnosing and monitoring ADHD in children” (Hall et al., 2016). This review identified and investigated six commercially available continuous performance test (CPT) used for aiding the diagnosis of ADHD, and two measures for objectively assessing activity in ADHD. The CPTs studied were the: Test of Variables of Attention (TOVA); Gordon Diagnostic System (GDS); Conners Continuous Performance Test (CCPT); Integrated Visual and Auditory Continuous Performance Test (IVA + CPT); Quotient ADHD system (or McLean Motion and Attention Test; MMAT) and the QbTest. The measures investigated for assessing activity in ADHD were based on: accelerometer-based devices (actigraphy and inertial measurement units (IMUs)) and infra-red motion analysis (MMAT and QbTest). From what is reported in this review, and can be considered comparable under similar conditions of testing, *prima facie,* the results obtained with the method advocated here, seems to be an improvement in accuracy, sensitivity and specificity over any of the objective tests scrutinised in the aforementioned systematic review.

Moreover to complement and partially update the former review, an *ad-hoc* in house “scoping review” was conducted. This review included other existing methods proposed to diagnose ADHD, that were not covered in the former systematic review. To make the indirect comparison possible, this in-house scoping review included only diagnostic accuracy studies that reported the chosen metrics of diagnostic for methods based on; electroencephalography and eventrelated potentials (Marcano et al., 2017; Loo et al., 2016; Snyder et al., 2015; Mohammadi et al., 2016; Gloss et al., 2016; Biederman et al., 2017; Gamma and Kara, 2016; Marcano et al., 2018; Manouilenko et al., 2017), structural and functional neuroimaging (Iannaccone et al., 2015; Rangarajan et al., 2014; de Celis Alonso et al., 2017; Qureshi et al., 2017; Serrallach et al., 2016; Hasaneen et al., 2017; Tan et al., 2017b; Uddin et al., 2017), simulated virtual reality and computer games (Negut et al., 2017, 2016; Berger et al., 2017; Faraone et al., 2016), and peripheral biochemical markers (Faraone et al., 2014; Scassellati and Bonvicini, 2015; Scassellati et al., 2012; Thome et al., 2012). According with this complementary “scoping review”, the method presented in this research, seems to outperforms all the diagnostic accuracy metrics reported in the trials scrutinised in the aforementioned review.

To conclude the discussion of this point, it is worthwhile to mention that this is the first study, which is based on objective measures that report a perfect diagnostic classification accuracy in a “training” and in an “independent” testing dataset of those with and without ADHD.

From a different vantage point, the method advocated by this research presents certain kind of advantages (over other alternative tests) that are related with its simplicity and appealing character (being a computer board game makes the task particularly attractive for children). Besides, its administration is straightforward and does not present major difficulties for children as little as five years old, to be handled, or requires any complex training of the tester. All these features are highly desirable for a test in a clinic situation.

An additional strength of the present study is its adherence to the STARD guidelines in reporting the classification performance of the models trialled (Bossuyt et al., 2015).

### 4.1 Study Limitations

In general terms, in a phase I and II of a diagnostic accuracy study, the limitations are directly related with the scope of the questions posed, and the adequacy of the methods employed to answer them (Knottnerus and Buntinx, 2008). Both, questions and methods set the boundaries to interpret the results.

In this regard, special care must be taken when attempting to go beyond those boundaries; most of the time, trespassing beyond them, leads to spurious generalisations. Besides, it is important to remark the proof-of-concept, exploratory nature of these studies and that they meant to be a first approximation to a new technology’s diagnostic value (Zhou et al., 2011; Larner, 2015; Dennis et al., 2009). After these former digressions, it is worthwhile to identify some of the sources of particular limitations that affected the present research.

The most relevant limitation is that the index test procedure was not performed in subjects with diverse manifestation and severity of ADHD or alternative similar diagnosis and comorbidities. The testing was restricted to children with ADHD-combined presentation and typically developing control children.

Another limitation of this study is that in a few cases, the determination of the reference standard (due its inherent subjective nature) presented uncertainties in identifying the true state (ADHD or non-ADHD) of the subject. Notwithstanding, since no major discrepancies existed between the reference standard and the index test, no measures were taken to control for the use of an imperfect gold standard (Hawkins et al., 2001).

Furthermore, due to the limited sample of ADHD cases and typically developing control children, the heterogeneity of behaviour in both groups was restricted. This lack of heterogeneity, despite that an independent testing dataset was used to evaluate the classification accuracy of the models, could have affected the classification outcome producing overly optimistic results.

### 4.2 Implications for Practise

Despite the encouraging results obtained in this research, it must be clearly stated that the method presented, in is actual form, is not ready to be used in a routine clinical scenario. Further translational development is mandatory, i.e., a standard, integrated version of the computer game and the analysis procedures should be produced. The computer application needs to be user-friendly and versatile to serve different clinical and research scenarios. It must contain a main module with the game and accessory modules to record, manage, perform data preparation/analysis and report the results. For the advance of the translational process it is highly recommendable that this research could be replicated by a different research team. After that phases III and IV diagnostic of accuracy studies should be performed.

## 5 Conclusions

The outcomes of this clinical research trial strongly support the present line of enquiry. There is a robust trend in the results, that provided direct evidence for the working hypotheses of this research, i.e:

i. The mean of the multiscale-measures from the ADHD group are different from the non-ADHD control group at the 0.05 significance level.

ii. Using a set of fractal measures (multifractal, lacunarity and multiscale straightness index), in a classification model, aimed to identify ADHD from non-ADHD cases, the area under the receiver operating characteristics ROC curve (AUC) is θ ϑ 0.80.

After performing indirect comparison but considering similar experimental conditions of testing, in a diagnostic accuracy scenario, these results seem, *prima facie*, to be extremely encouraging, making a compelling case for further investigation of the suitability of this novel approach when compared to current clinical objective tests used to assist in the diagnosis of ADHD.

From a translational vantage point, the technology presented here makes it a promising candidate to further develop a screening, diagnostic and monitoring system for ADHD(Faraone et al., 2014; Thome et al., 2012), due to its straightforward application, simple-to-perform, reliable, reproducible, inexpensive and non-invasive nature. Another asset of this technology is that potentially could be easily adapted to assess other EF disorders. Nevertheless, before any clinical inception of the testing procedures advocated by this research; replication studies should be performed, and phases III and IV of diagnostic research’s questions should be answered.

Summing up, these results strongly support the hypotheses contended in this research and show that targeting the dynamics of human inferential processes is a promising way to deal with EF disorders in a diagnosis scenario.

## 6 Other Information

### Acknowledgement

The authors gratefully acknowledges and gives special thanks and gratitude to all the staff from the Paediatrics Department at the Nelson Marlborough District Health Board and the Iwi Health board for supporting this clinical trial. In addition, we wish to thank to all the children and families from the Nelson community that participated in the trial.

## Footnotes

1 In this respect, Galen’s opinion that, *“so long as we are ignorant of the true essence of the cause which is operating, we call it a faculty*”, expresses very much what is contended here (Galen, trans.1916).

